# Engineering Modular and Tunable Single Molecule Sensors by Decoupling Sensing from Signal Output

**DOI:** 10.1101/2023.11.06.565795

**Authors:** Lennart Grabenhorst, Martina Pfeiffer, Thea Schinkel, Mirjam Kümmerlin, Jasmin B. Maglic, Gereon A. Brüggenthies, Florian Selbach, Alexander T. Murr, Philip Tinnefeld, Viktorija Glembockyte

**Author notes:** Correspondence should be addressed to P.T. and V.G. These authors contributed equally (Lennart Grabenhorst, Martina Pfeiffer).

## Abstract

Biosensors play key roles in medical research and diagnostics, but there currently is a lack of sensing platforms that combine easy adaptation to new targets, strategies to tune the response window to relevant analyte concentration ranges and allow for the incorporation of multiple sensing elements to benefit from multivalency. Utilizing a DNA origami nanostructure as a scaffold for arranging the different sensor components, we here propose an approach for the development of modular and tunable single-molecule sensors capable of detecting a variety of biomolecular targets such as nucleic acids, antibodies and restriction enzymes while offering mechanisms to tune the dynamic window, the specificity, and the cooperativity of the sensor.

---

Fluorescent sensors are our gateway to a deeper understanding of cellular processes [1–4] and diseases [5–7]. A typical biosensor consists of two functional units: a biorecognition element capable of sensing an analyte or biological activity and a signal transduction element translating it into measurable readout. For virtually every biomolecular target of interest to medical research or diagnostics, it is possible to find a molecule (e.g., antibody, receptor, or aptamer) that binds to it with high specificity and sensitivity. Given that the conformational change upon target binding is often very small, one of the key challenges in transforming these binders into useful fluorescence sensors lies in achieving a measurable fluorescence signal (e.g., change in fluorescence intensity or Fluorescence Resonance Energy Transfer (FRET) between donor and acceptor labels) upon binding. This challenge has been addressed by a number of elegant modular strategies which generalize and simplify the development of new biosensors, for example, by engineering superstructures that elicit large conformational changes upon target binding with the help of semi-synthetic protein chimeras [8–13], chemically induced dimerization [14, 15], de novo protein design [15, 16] or by using conditionally stable ligand-binding domains [17, 18].

A second fundamental challenge in developing new sensors lies in tailoring their response window to the analyte concentration of interest. Binding of a ligand to a single-site recognition element produces a hyperbolic dose-response curve with a fixed response window spanning roughly two orders of magnitude. This limits the utility of the sensor in applications that require either great sensitivity (sharp signal response) or quantification of target molecules in a concentration window that varies or spans several orders of magnitude [19]. Mimicking nature’s tricks to overcome this challenge, mechanisms of allosteric control [20–23], sequestration [24], and cooperativity [25–29] have been implemented in synthetic sensor and signaling systems. While these approaches have demonstrated impressive tuning capabilities in aptamer-based sensors enabling measurement of target concentrations across orders of magnitude [22] or narrowing of the response window to as little as 3-fold, they lack the modularity required for straightforward extension to arbitrary analytes. Possible ways to shift the response window of the sensor (e.g., by introducing a mutation in the binding site [11, 14, 15, 17] or changes to the scaffolding structure [8, 9, 16]) have also been outlined in modular sensing platforms, yet, most of these approaches rely on single-site binding and cannot harness additional tuning and design advantages available to multivalent sensors (e.g., cooperativity or multiplexing).

Strategies to simultaneously decouple sensing from signal transduction, tune the response window of the sensor and combine multiple sensing elements are of great interest, as they would allow independent tuning of sensor properties and thus greatly increase the speed at which new sensors can be developed. However, a global sensor approach that addresses all these challenges in one platform has yet to be realized. To lay out fundamental strategies that could combine all these requirements, in this work we harnessed the nanoscale arranging capabilities and modularity of DNA origami nanostructures. Using a dynamic nanostructure to assemble different sensor elements, we obtained almost digital FRET signal readout with single-molecule sensitivity, outlined strategies to tune the response and specificity of a sensor as well as developed multiplexed sensors capable of more complex sensing operations.

## Engineering of a spatially decoupled signal transduction element

To decouple sensing from signal output and build a sensor platform with a high optical contrast, we utilized a dynamic DNA origami nanostructure [30–32] capable of undergoing large conformational changes. It consists of two ca. 65-nm long arms connected by a single-stranded (ss) scaffold DNA region (Fig. 1a). In absence of additional interactions, the two arms fluctuate around an equilibrium angle of ca. 90° (Fig. 1b, upper panel). However, by introducing DNA-DNA closing interactions on the two arms it can be folded and purified in a closed conformation, with both arms almost parallel to each other (Fig. 1b, lower panel). In the model nanosensor, closing interactions are designed with a ssDNA overhang allowing for toehold-mediated opening of the structure by complementary ssDNA opening strands. At the same time, a short toehold overhang is left upon binding the target, enabling reversible reclosing and mimicking receptor-ligand interactions (Figs. 1a, S1, S2, and S16).

**Figure 1:**
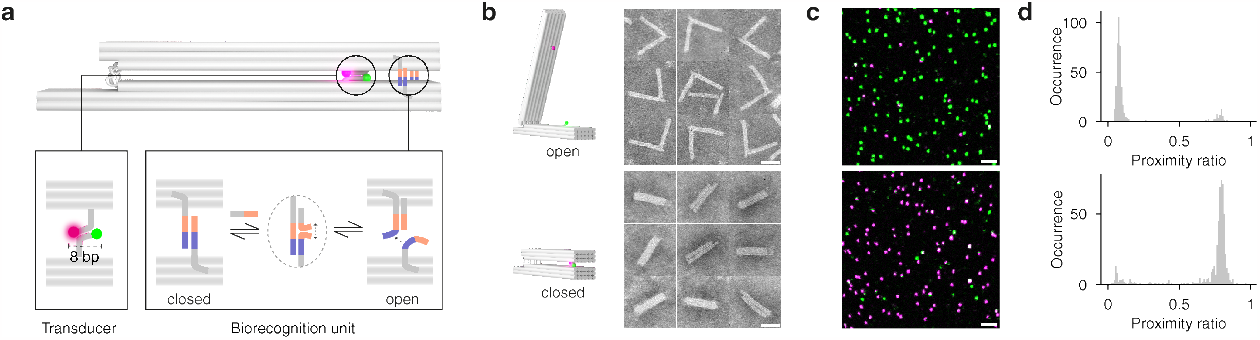
Design of the modular biosensor platform. **(a)** DNA origami nanostructure used to arrange and decouple sensing from signal output. The signal transduction element consists of a donor (ATTO542) and an acceptor (ATTO647N) dye forming a FRET pair (bottom left) brought together in the orientation required for high FRET in the closed state by weak 8 base pair (bp) DNA hybridization. In the model sensor platform, the biorecognition element is mimicked by a reversible closing interaction based on toehold-mediated DNA strand displacement reaction(s) (bottom right). **(b)** Snapshots from transmission electron micrographs of the sensors in each of the conformations (scale bar: 40 nm, for full micrographs see Fig. S3). **(c)** Confocal microscopy scans of surface immobilized biosensors in the open (top) and closed (bottom) conformation (scale bar: 2 µm). **(d)** Corresponding spot wise PR histograms illustrating the high FRET contrast between closed and open states of the sensor.

The signal transduction element consists of bright and photostable donor (ATTO542) and acceptor (ATTO647N) fluorophores positioned on the opposing arms forming a FRET pair (Fig. 1a, lower left). We studied different sensor designs at the single-molecule level by incorporating biotinylated ssDNA strands to immobilize the structures on BSA-biotin/Neutravidin-coated coverslips and performing confocal scans. The extent of FRET in different sensor designs was characterized by calculating the proximity ratio (PR, defined as I_Red_/(I_Red_ + I_Green_)) for each nanosensor. The signal output was optimized to minimize the effect of the flexibility of the nanostructure and the impact of changes in ionic strength on the PR distributions (Fig. S4, oxDNA [33] simulations in Videos S1-S2). To this end, two complementary ssDNAs protruding from each of the arms were used to define the positions of the two dyes in the closed state, leading to a narrow distribution of high FRET values (⟨ PR_closed_⟩ ca. 0.77, Fig. 1c and 1d, top). The large conformational change upon opening the sensor with ssDNA target separates the dyes in the FRET pair resulting in negligible FRET in the open state (⟨ PR_open_⟩ ca. 0.08, Fig. 1d, bottom). This high optical contrast allowed for the unambiguous assignment of the two conformational states and enabled the detailed characterization of the sensor response.

### Tuning the response window of the biosensor

A universal sensor platform would allow to readily assemble new sensors from a large pool of already existing receptor-ligand interactions. This calls for strategies to tune both, onset, and sharpness, of the sensor response to enable monitoring relevant concentration changes of target molecules without the need to re-engineer these interactions. In natural receptor-ligand systems, this is commonly achieved by hierarchical assembly of multiple binding units or allosteric modulation of the binding interaction [34]. We laid out and tested several strategies inspired by these mechanisms to tune the signal response of the nanosensor without altering the sensing interaction itself. The high signal contrast between open and closed conformations enabled us to read out the equilibrium distributions with high precision and thus conduct single-molecule titration experiments with the ssDNA target as a model ligand (Fig. 2a). By quantifying the fraction of open sensors (PR<0.3, Fig. 2b) at each target concentration and fitting the resulting binding curve with the Hill equation, we characterized the response window of each sensor design in terms of overall affinity K_½_ which represents the target concentration where half of the sensors are open, and the Hill coefficient n_H_, which is a measure of the cooperativity in a system (Fig. 2b).

**Figure 2:**
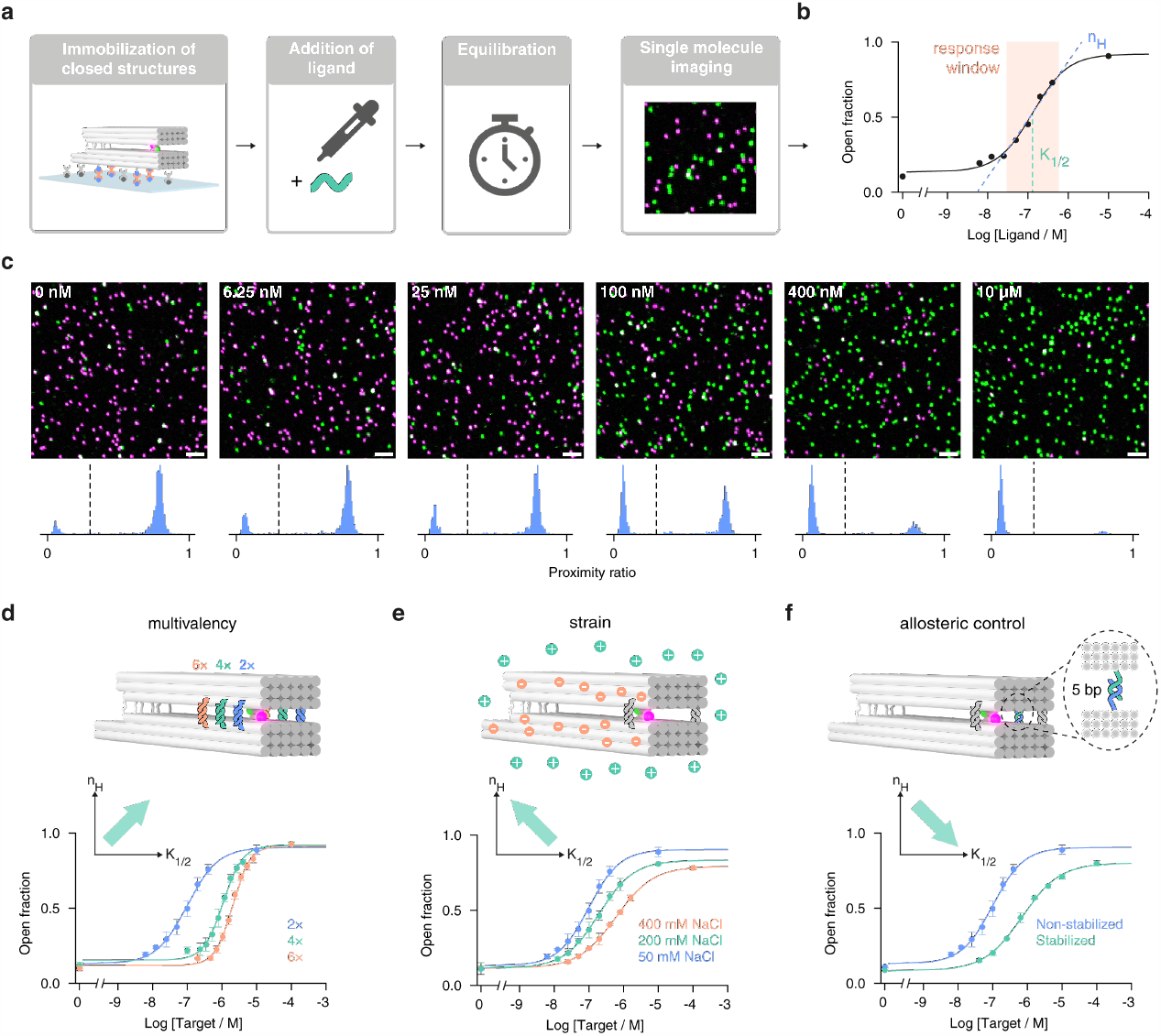
Studying and tuning the response window of the sensors on the single molecule level. **(a)** Workflow to investigate the response window of different sensor designs: sensors are immobilized via Biotin-NeutrAvidin interactions. For each target concentration, one surface is prepared, and a defined concentration of target is added. After equilibration, confocal scans are acquired, and the fraction of closed structures is determined by analyzing on average 681 single nanosensors for each concentration. **(b)** The response window, i.e., the overall affinity of the sensor (K_½_) and the extent of cooperativity (n_H_) are determined by fitting the titration curve with the Hill equation. **(c)** Example confocal fluorescence scans for different target concentrations with respective PR histograms shown below (scale bar: 2 µm, dashed line represents the PR threshold used to assign sensors as open). **(d)** Increasing the number of sensing elements shifts the K_½_ of the sensor to higher values and increases the cooperativity. **(e)** Decreasing the ionic strength increases the coulombic repulsion and destabilizes the closed state of the sensor, leading to lower K_½_ and higher n_H_. **(f)** Introducing additional DNA-DNA stabilization interactions stabilizes the closed state of the sensor and leads to an increase of K_½_ and a decrease of n_H_. Titration curves show mean values of three independent measurements with the error bars corresponding to the standard deviation.

Inspired by naturally occurring multivalent systems, we rationalized that increasing the number of sensing interactions would provide means to engineer cooperativity in our nanosensors. As expected, multivalency induced cooperative behavior to our system: going from two to four interactions, n_H_ increased from 0.98±0.08 to 1.55±0.15, respectively, while in the sensor with six interactions, we achieved a cooperativity of 1.73±0.13. Additionally, going from two to four sensing elements led to an almost 10-fold increase in K_½_ (from 100±10 nM to 1.09±0.09 µM, respectively) and adding two more closing interactions further doubled K_½_ to 2.03±0.10 µM (Fig. 2d). We propose that this cooperativity is a result of strain in the closed state of the sensor acting on all interactions induced by coulombic repulsion (i.e., opening of one interaction increases the force acting on the remaining ones). Altogether, increasing the number of sensing elements in these multivalent sensors provided a strategy to sharpen the signal response (narrowing the response window from ≈80-fold to ≈12-fold) due to arising cooperativity while simultaneously shifting K_½_ to higher values (Fig. 2d, inset).

To introduce an orthogonal tuning strategy, we set out to alter the force that the backbone structure exerts on the closing interactions. We reasoned that this can be achieved by increasing coulombic repulsion in the closed state [31, 32] and, in turn, characterized the properties of the sensor containing two closing interactions at varying ionic strengths (Fig. 2e). As expected, decreasing the NaCl concentration from 400 mM NaCl to 50 mM led to an earlier response, shifting K_½_ by 6-fold from 601±20 nM to 100±10 nM, which we attributed to destabilization of the closed state of the sensor. Interestingly, the cooperativity was also sensitive to the ionic strength: we obtained n_H_ values of 0.81±0.02, 0.87±0.04, 0.98±0.08 in the presence of 400 mM, 200 mM, and 50 mM NaCl, respectively (Fig. 2d). This result is in line with our earlier assumption that cooperativity increases with coulombic repulsion. The negative cooperativity obtained at higher ionic strengths, however, highlights the existence of a competing cooperative process. Here, multiple closing interactions facilitate the reclosing, in turn, providing means to extend the response window of the nanosensors. Overall, the destabilization of the closed state provided a mechanism to increase the cooperativity and shift the response window to lower concentrations providing an orthogonal direction in the n_H_ vs. K_½_ space (insets in Fig. 2d and 2e).

Finally, by including additional ssDNA strands that hybridize to each other in the closed conformation we investigated the possibility to implement allosteric control. Introduction of two transient DNA-DNA interactions on the inside of the two arms of the structure led to a 7-fold increase of K_½_ from 100±10 nM to 705±65 nM (Fig. 2f) illustrating the sensitivity of this platform to small alterations. Varying the strength of these stabilizing interactions provides a mechanism to fine-tune the overall affinity of the nanosensor. Analogous to what was observed at increasing ionic strengths (Fig. 2e), the stabilization of the closed state via this strategy has led to a decrease in n_H_ from 0.98±0.08 to 0.8±0.05 consistent with facilitated reclosing. Altogether, the three illustrated approaches present strategies to tune the onset (K_½_) and sharpness (n_H_) of the sensor without altering the sensing interaction itself with strategies that are orthogonal to each other (insets in Figs. 2d-f) and can be combined to cover an extended K_½_ and n_H_ parameter space, something that so far has been challenging to implement in synthetic sensor systems [27, 28].

### Harnessing multivalency for increased target specificity

One of the key properties sought after in biosensors is the ability to detect target analytes specifically in a large pool of other similar biomolecules. As such, we investigated whether multivalency could be utilized to improve the specificity of the nanosensor. We rationalized that increasing the number of sensing elements would amplify the overall binding energy difference of two energetically similar targets. To confirm this, we first studied the opening of a nanosensor containing four sensing interactions in the presence of 17-nt perfectly matching target as well as targets containing one (C-C mismatch, ΔΔG from the perfectly matched target of 5.4 kcal/mol as estimated by NUPACK [35], Fig. S5) and two (C-C + T-T mismatch, ΔΔG 6.2 kcal/mol) nucleotide mismatches (Fig. 3a). For the perfectly matched target the nanosensor opened at nanomolar concentrations, whereas in the presence of targets containing one or two nucleotide mismatches no opening was observed even at 10 µM (Fig. 3b) illustrating the ability of the nanosensor to specifically detect perfectly matched targets even in ≈ 1000-fold excess of similar off-targets.

**Figure 3:**
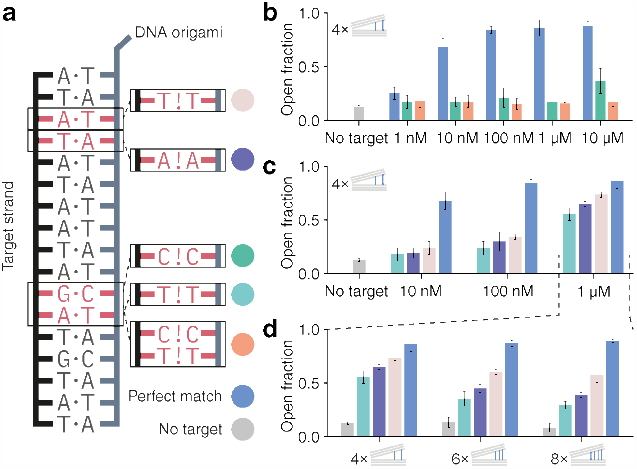
Harnessing sensor multivalency for increased target specificity. **(a)** Schematic of the model receptor-ligand opening interaction (left). For testing specificity of different sensor designs, we included mismatches into the opening interaction (right), mimicking a ligand (off-target) with similar binding strength. **(b)** Comparing the perfectly matched opening interaction (blue) to an interaction with one (green) and two (orange) single nucleotide mismatches shows drastic differences in sensor response: while perfectly matched target (blue) opens the sensor at nanomolar concentrations, both mismatched targets exhibit almost no response up to 10 µM concentration. **(c)** Testing targets with even smaller free energy differences from the perfect match. **(d)** Increasing the number of closing interactions from 4 to 8 increases the capabilities of the sensor to differentiate these targets from the perfect match as illustrated with the sensor opening measured in the presence of 1 µM target concentrations. Plots show the mean values of three independent measurements with the error bars corresponding to the standard deviation. Examples for all confocal fluorescence scans are found in Fig. S6.

Exploring this potential further, we investigated whether we could differentiate off-targets with even smaller ΔΔGs (2.3 — 3.6 kcal/mol, A-A and T-T mismatches, Fig. 3c and Fig. S5). By incubating the nanosensor containing four sensing interactions with different targets, we could show that at up to 100 nM target concentration it is possible to clearly differentiate the energetically similar off-targets by both comparing the equilibrium fraction of open nanosensors or by monitoring the nanosensor opening kinetics (Fig. S7). Nonetheless, as the concentration of the target strands is further increased to 1 µM (Fig. 3c) this differentiation becomes more difficult. While the specificity of the nanosensor at micromolar concentrations of targets may be less relevant for detection of nucleic acids, we deemed that it is still crucial for the utility of the modular sensing platform, where the desired response of the sensor lies at higher concentrations (e.g., sensing of metabolites). We conducted additional studies with nanosensors containing 6 and 8 closing interactions to demonstrate that multivalency can further improve the specificity when desired. As shown in Fig. 3d, increasing the number of sensing interactions increased the differences in opening fraction when compared to the perfect match. With the nanosensor containing 8 closing interactions, we were able to differentiate between two off-targets with single T-T and A-A mismatches that have an estimated difference in binding energy as little as 0.51 kcal/mol (less than 1 k_B_T, Fig. S5).

### Extension to other biomolecular targets and incorporation of logic sensing operations

We next demonstrated that the large optical signal contrast developed in our model nanosensor can indeed be swiftly and modularly adapted to other biomolecular targets without the need to tediously re-optimize the signal output. A single molecule sensor for anti-Digoxigenin (anti-Dig) antibodies was engineered by incorporating two specific Dig antigen biorecognition elements on each sensor arm (Fig. 4a). To meet the required geometry for bivalent binding36 the nanosensor was initially kept in the closed state via DNA-DNA interactions. After surface immobilization and antibody incubation, DNA opening staples were added which resulted in ≈ 85% sensors still in the closed state (Fig. 4a, upper panel and Fig. 4b) suggesting that the sensor is held in the closed state by the bivalent binding of the antibody. However, in the absence of the antibody, the nanosensors opened almost quantitatively (Fig. 4a, lower panel and Fig. 4b). As illustrated in Fig. 4a and 4b, the near digital signal contrast optimized for the model sensor is still preserved in the antibody nanosensor. Additionally, we evaluated the specificity (Figs. S8 and S9) as well as the potential of this antibody sensor to be useful in more complex biological fluids by performing the antibody assay described above in 50% blood plasma (Fig. 4c and Fig. S10). The obtained percentages of open sensors measured in blood plasma were within experimental error when compared to those measured in buffer, confirming that neither stability of the DNA origami nanosensor nor the performance of the antibody binding assay were compromised. We also evaluated the sensitive concentration range of the anti-Dig antibody assay under clinically relevant conditions (20 min incubation) to show that the antibody concentration at which the signal change is half the maximum (C_half−max_) lies at 104 pM, which is in accordance to previously reported values [37] and in a concentration range relevant for diagnostic applications [38] (Fig. S11).

**Figure 4:**
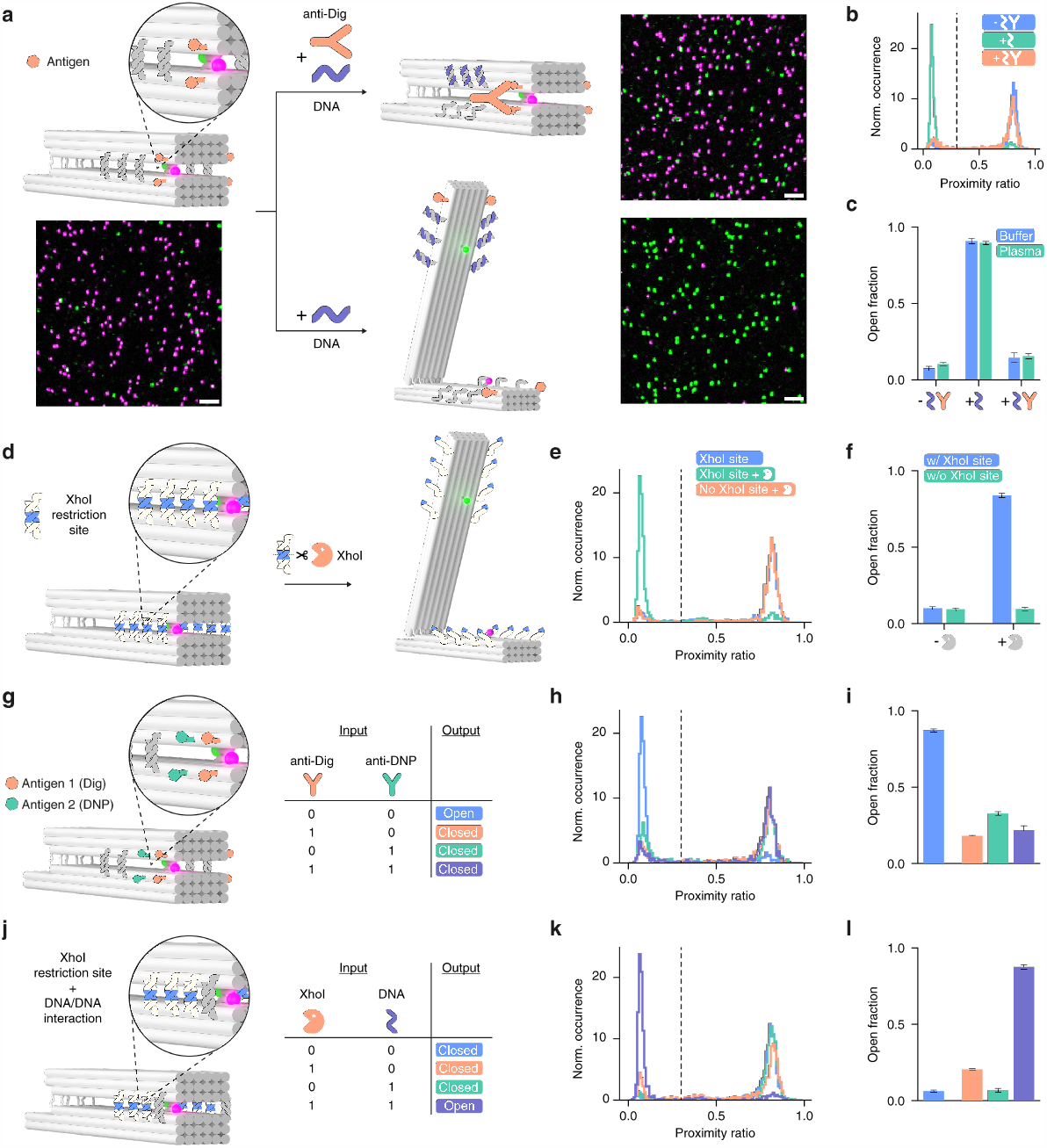
Extension of the sensor platform to different molecular targets and introduction of logic sensing operations. **(a)** Detection of antibodies by incorporation of antigen recognition elements (Dig) on the opposing arms of nanosensor kept in the closed state via DNA-DNA interactions. Upon addition of DNA opening strands the nanosensor stays closed in the presence and opens in the absence of anti-Dig antibodies. The high signal contrast is illustrated by confocal microscopy scans (scale bar: 2 µm). **(b)** Spot-wise PR histograms obtained for the antibody sensor in the absence (blue) and in the presence of DNA opening strands (cyan) as well as in the presence of DNA opening strands and 100 nM anti-Dig antibody (orange). **(c)** Corresponding fraction of open antibody nanosensors obtained in buffer as well as in blood plasma. **(d)** Detection of enzymatic activity by incorporation of nuclease-specific restriction site in the DNA-DNA closing interaction. Spot wise PR histograms obtained for nanosensors containing the XhoI restriction site in the absence (green) and presence (blue) of XhoI. No opening was observed for nanosensors without the specific restriction site in the presence of XhoI (orange). **(f)** Corresponding fraction of open nuclease nanosensors with and without restriction site. **(g)** Multiplexed detection of two different antibodies (anti-Dig and anti-DNP) via a molecular OR gate is achieved via the incorporation of two different antigens (Dig and DNP) on the opposing arms of the closed nanosensor. Upon addition of DNA opening strands the nanosensor stays closed if either or both antibodies are present. **(h)** Spot wise PR histograms for antibody nanosensor inputs shown in (g). **(i)** Corresponding fraction of open nanosensors for different antibody nanosensor inputs. **(j)** Multiplexed detection of two different biomolecular targets (nucleic acid and restriction enzyme) via a logic AND gate can be achieved by incorporation of the restriction site and another DNA-DNA closing interaction. The nanosensor opens only in the presence of both, nucleic acid, and restriction enzyme targets. **(k)** Corresponding spot wise PR histograms obtained for different molecular inputs of shown in (j). **(l)** Corresponding fraction of open nanosensors for different nanosensor inputs. Bar plots show the mean values of three independent measurements (each include at least 423 single nanosensors) with the error bars corresponding to the standard deviation.

Next, we studied whether the sensor platform can also be utilized for the detection of enzymatic activities. Usually, optical enzyme (e.g., protease or nuclease) activity sensors are designed by placing two labels close to the substrate binding site in a manner that leads to loss of FRET or turn-on of fluorescence signal upon substrate cleavage. In designing such sensors, one is faced with the inherent conflict between placing the labels close enough to the cleavage site to achieve high optical contrast yet far enough not to influence the enzyme-substrate binding which results in many rounds of optimization and often suboptimal signal contrast. This challenge can be globally addressed by the sensor scheme proposed here where target recognition is decoupled and spatially separated from the transduction element. To illustrate this, we designed a single molecule activity sensor for the nuclease XhoI: without the need to re-engineer the transduction element, we simply incorporated the restriction sites for XhoI in the DNA-DNA closing interactions (Fig. 4d). In the presence of XhoI, the closing interactions are cleaved leading to opening of the DNA origami sensor and loss of FRET (Figs. 4e and S12). In contrast, no opening is observed if the nanosensors are closed with DNA-DNA interactions without XhoI cleavage sites, confirming the desired specificity (Figs. 4e, 4f, and S13).

Finally, to illustrate the ability of the proposed nanosensor to modularly exchange and combine different sensing elements on one platform, we designed a sensor capable of detecting two different (anti-Dig and anti-DNP) antibodies (Fig. 4g) acting as a molecular OR gate. Two different antigens (Dig and DNP) were incorporated on the opposing arms of the nanosensor closed via DNA-DNA interactions. Upon addition of DNA opening strands the nanosensor opens when neither of the antibodies are present and stays closed if either or both antibodies are present (Fig. 4i). Fig. 4h and Fig. S14 illustrate that the high FRET contrast is still preserved despite a more complex sensing scheme. In fact, as illustrated in Fig. 4j, the multiplexed sensing is not restricted to the same type of biorecognition elements: by incorporating the XhoI restriction site and another DNA-DNA closing interaction we could build a logic AND gate for two different biomolecular targets: one based on a binding event, and one based on a cleavage reaction. Here we could show that the sensor only opens when both, XhoI and DNA targets, are present (Figs. 4k, l, and Fig. S15). Altogether, the modularity the DNA origami offers to incorporate different biorecognition elements, combined with the preserved robust FRET readout opens many possibilities to engineer new multiplexed sensors as well as logical sensing schemes for answering more complex diagnostic questions.

## Conclusions

General and modular strategies to assemble sensors have the potential to vastly speed up the development of new diagnostic tools for health and disease research. Here, we developed a generalizable approach to create DNA origami-based synthetic sensing systems with high signal contrast and single-molecule sensitivity. We harnessed the modularity of DNA origami nanostructures to: 1) modularly assemble all the elements of the sensor, 2) spatially decouple sensing from signal output, 3) provide a large conformational change required for large FRET contrast, 4) implement strategies to tune the response window, 5) engineer multivalent sensors which enabled improved specificity, multiplexing, logic sensing, and additional tuning capabilities via cooperativity.

Optical single molecule sensors allow to detect and monitor target analytes with ultimate sensitivity and unprecedented spatial resolution. However, achieving a large FRET contrast in single-molecule DNA origami sensors so far has been challenging: low FRET contrast and broad FRET distributions often require averaging over hundreds of nanostructures to distinguish the two states of the sensor [39–42]. Here we combined two tricks to solve this: a large conformational change and an additional transient guiding interaction to control the orientation of the dyes in the FRET pair – which led to a high FRET contrast and allowed the clear differentiation of the two sensor states on the single molecule level. The modularity and the ability to immobilize the sensors on the surface in a specific orientation [43] make this approach readily extendable to other readout mechanisms, such as electrochemical readout [44], fluorescence quenching (e.g., by a graphene surface [45, 46]), or bioluminescence energy transfer [9, 11, 17] as well as pave the way towards highly multiplexed sensor chips.

An exciting avenue to explore going forward would be the utilization of de novo designed binders [15, 16] as sensing elements, combining the many possibilities provided by protein design with a high-contrast single molecule readout and multivalent sensing schemes. One of the challenges that would have to be addressed, however, is the stability of DNA origami-based sensors. Here we showed that the sensor is still functional in 50% blood plasma, but further studies will be needed to fully assess the stability of the sensors in complex chemical environments or live cells. Recent progress on stabilization strategies for DNA origami structures [47–49] offers many options that can be tested to achieve maximal performance. Altogether, the modularity, tunability, and sensitivity of the reported approach provide a starting point for the rapid development of tailored and complex sensors for a wide range of analytes.

## Acknowledgements

We thank Prof. T. Liedl and Prof. J. Rädler for providing access to transmission electron microscopy facilities and the members of the Tinnefeld lab for discussion and feedback, especially Dr. F. Steiner. We also thank Michael Scheckenbach for initial AFM measurements.

## Author contributions

V.G. and P.T. conceived the idea and directed the project. V.G., P.T., L.G. and M.P. further conceptualized the research. L.G. designed the DNA origami sensor and performed oxDNA simulations. L.G., V.G., M.K., and J.M. tested and optimized the design of the sensor and the signal transduction element. F.S. and G.A.B. performed TEM measurements. L.G. and V.G. implemented the sensor tuning strategies and carried out and analyzed single molecule titration experiments with the help of G.A.B. and A.T.M.; T.S. and V.G. designed and carried out the sensor specificity studies. M.P. carried out and analyzed antibody, nuclease detection, multiplexed detection assays. L.G. wrote the code for single-molecule data analysis. L.G. and V.G. prepared figures. V.G., L.G. and P.T. wrote the manuscript with additional input from M.P.; all authors reviewed and accepted the manuscript.

## Data and materials availability

All experimental data supporting the findings of this work as well as DNA origami cadnano files will be made available from a public repository. The custom code for the analysis of single molecule confocal scans is available online (https://gitlab.lrz.de/tinnefeldlab/cospota).

## Methods

### DNA origami design and synthesis

DNA origami nanostructures were designed in caDNAno [1] according to a design published by Marras et al. [2]. All staple strands were ordered at Integrated DNA Technologies, Inc., Belgium except for the fluorescently labeled strands, which were ordered at biomers.net GmbH, Germany. For detailed folding recipes and sequences, see Supplementary Text and Tables S1-S8. The folding was executed in a thermocycler using the temperature ramp as described earlier [3]. After folding, samples were subjected to agarose gel electrophoresis; we employed 1.5% w/v gels in 1×TAE buffer supplemented with 10 mM MgCl_2_. Gels were run for at least 3 hours at 70 V. Then, gels were inspected using a gel documentation system (Fusion FX, Vilber Deutschland GmbH, Germany) where we used the red fluorescence channel to identify the bands containing closed DNA origami sensors (an example scan is shown in Fig. S16). These bands were cut using a scalpel and placed on Parafilm. Then, the gel fragments were squeezed with a small glass slide wrapped in Parafilm to extract the DNA origami nanostructures. For storage, we aliquoted the origami solution to 200 µL PCR tubes: we mixed 40 µL of origami solution with 8 µL of 5 M NaCl solution to minimize degradation. Two slightly different versions of the nanosensors were used for different applications - for details on the differences, see Tables S1-S7 and supporting text.

### Coarse-grained simulations of DNA origami nanostructures

caDNAno design files were converted to oxDNA [4–6] input files using the tacoxDNA web server [7], the closing interactions were manually added in oxView [8] and the resulting structure was simulated at 20 °C, 400 mM NaCl on the oxDNA.org server [9] with the standard settings as well as the recommended relaxation procedure.

### Transmision electron microscopy (TEM) measurements

TEM grids (Formvar/carbon, 400 mesh, Cu, TedPella, Inc., USA) were Ar-plasma cleaned and incubated for 2 min with DNA origami sample (5 µL, 1-5 nM). Grids were washed with 2% uranyl formate solution (5 µL) and incubated for another 4 s with 2% uranyl formate solution (5 µL) for staining. TEM imaging was performed on a JEM-1100 microscope (JEOL GmbH, Japan) with an acceleration voltage of 80 kV.

### Preparation of microscopy samples

Microscope slides (24 mm × 60 mm size and 170 μm thickness (Carl Roth GmbH, Germany)) were cleaned in a UV/ozone cleaner (PSD-UV4, Novascan Technologies, USA) for 30 mins at 100 °C. Then, CoverWell perfusion chambers (Grace Bio-Labs, 0.5 mm deep) were glued on top of the slides and the glue was strengthened by heating on a hot plate for ca. 20 s at 100 °C. Then, the chambers were cleaned with 1 M KOH by incubating for 10 mins and then washing with 1× PBS four times. Surfaces were passivated by incubating with BSA-Biotin (1 mg/mL in 1× TE with 50 mM NaCl, Thermo Fisher Scientific, USA) and subsequent washing with 1× PBS buffer three times. Then, chambers were incubated with 0.25 mg/mL NeutrAvidin (Thermo Fisher Scientific, USA) in 1× PBS for 10 mins followed by final washing with 1× PBS three times. DNA origami solutions were diluted in immobilization buffer (10 mM TRIS, 10 mM MgCl_2_, 750 mM NaCl) as required to reach the desired surface density of DNA origami sensors (50–100 pM), which was checked on the confocal microscope. When the desired density was reached, the surfaces were washed in the respective buffer that was used for the experiment.

### Single-molecule confocal microscopy measurements

Home-built confocal microscope based on an Olympus IX-83 body (Japan) was used to acquire single molecule fluorescence scans. A supercontinuum white light laser pulsed at 78 MHz (SuperK Extreme, NKT Photonics, Denmark) was used to excite the samples. The excitation wavelength was selected using an acousto-optically tunable filter (SuperK Dual AOTF, NKT Photonics, Denmark) controlled by a digital controller (AODS 20160 8R, Crystal Technology, USA). If needed, a second acousto-optically tunable filter (AA.AOTF.ns:TN, AA Opto-Electronic, France) controlled with a home-written LabVIEW (National Instruments, USA) program was used to alternate between two wavelengths (532- and 639-nm). We used a neutral density filter, a linear polarizer and a λ/4 plate to achieve circularly polarized excitation light of the desired intensity. For the single-molecule confocal scans 532-nm (2 µW) excitation was used. The excitation light was sent to the sample through an oil immersion objective (UPlanSApo 100×, NA=1.4, WD=0.12 mm, Olympus, Japan) via a dichroic beam splitter (zt532/640rpc, Chroma, USA). A piezo stage (P-517.3CL, E-501.00, Physik Instrumente GmbH&Co. KG, Germany) was used to move the sample in X-Y. The sample was kept in focus over extended periods with the help of a z-drift compensation system (IX3-zdc2-83, Olympus, Japan). The emitted light was collected with the same objective and separated from the excitation light with the same dichroic beam splitter. Then, the emission light was sent through a 50-µm pinhole, the two emission channels were split using beam splitter (640DCXR, AHF Analysetechnik AG, Germany), additionally filtered with two emission filters (RazorEdge 647, Semrock Inc., USA for the red channel and BrightLine HC 582/75, Semrock Inc., USA for the green channel) and focused onto the single photon detectors (SPCM, AQR 14, PerkinElmer, USA). A custom LabVIEW program was used for data acquisition.

### Single-molecule wide-field microscopy measurements

For detection of single-molecule fluorescence transients, a commercial wide-field/TIRF microscope Nanoimager from Oxford Nanoimaging Ltd. was used. Red excitation at 638 nm was realized with a 1100 mW laser, green excitation at 532 nm with a 1000 mW laser, respectively. The relative laser intensities were set to ca. 3.5 mW for green and to ca. 1 mW for red excitation. The microscope was set to TIRF illumination. Data acquisition was initialized by time-lapsed imaging. A frame of 100 ms was recorded every second separately for both excitation lasers. Measurements were carried out at 37 °C.

### Single-molecule titrations

For each DNA target concentration a separate chamber with closed nanosensors (Version 2) was prepared. Single-molecule titrations were conducted in the buffer containing 10 mM Tris and 10 mM MgCl_2_ with varying amounts of NaCl (see Table S9 for details). Serial dilutions of the 15 bp opening strand (see Table S8) were prepared in the same buffer and each chamber was filled with the opening staple solution and closed using adhesive seal tabs (Grace Bio-Labs, USA). The samples were incubated overnight at room temperature to ensure that the thermal equilibrium has been reached even for very low (nM) target concentrations. The opening of the sensors was quantified by acquiring single-molecule fluorescence scans on the confocal microscope.

### Single-molecule pulsed interleaved excitation FRET measurements in solution

For single-molecule FRET measurements of freely diffusing nanostructures (Fig. S7) we used a home-built setup based on an Olympus IX-71 inverted microscope (for a detailed description of the setup, see Ref. [10]). The slides were prepared analogously to single-molecule measurements for surface immobilized samples except the slides were not incubated with NeutrAvidin to measure freely diffusing nanosensors in solution at 500 pM–1 nM concentrations. We used pulsed interleaved excitation with a 532-nm laser at 4 µW intensity and a 640-nm laser at 5 µW excitation intensity at 40 MHz repetition rate, respectively. The data was analyzed with the PAM [11] software. All channel burst search was performed to select the single bursts and ALEX 2CDE and |TDX-TAA| filters were used the filter the selected burst data.

### Single-molecule studies of target specificity

Nanosensors used for studying target specificity (Version 1) contained a shorter 11 bp closing interactions (for a full list of sequences see Table S8). For each (off) target concentration a separate chamber with closed nanosensors was prepared, filled with the solution containing the target DNA in 10 mM Tris, 1 mM EDTA, 5 mM MgCl_2_ and 750 mM NaCl and closed using adhesive seal tabs (Grace Bio-Labs, USA). Samples were incubated overnight to ensure that thermal equilibrium was reached, and the open fraction was determined by performing single-molecule fluorescence scans on the confocal microscope.

### Antibody detection assay

Nanosensors used for the detection of IgG antibodies (Version 1) were equipped with two digoxigenin (Dig) functionalities and/or two dinitrophenol (DNP) functionalities per sensor arm (see Table S7 for modified sequences). Additionally, four and six 11 bp DNA-DNA closing interactions, respectively, were incorporated to facilitate bivalent binding of the target antibody in a bridge-like manner to the opposite arms of the nanosensor (Table S3). For each sample a separate chamber with closed nanosensors was prepared. anti-Dig antibodies (Rb monoclonal, Thermo Fisher Scientific, USA, Catalog # 700772, AB_2532342) and/or anti-DNP antibodies (Rat monoclonal, Thermo Fisher Scientific, USA Catalog # 04-8300, AB_2532964) were diluted to 100 nM in the buffer or diluted heparin blood plasma containing 10 mM Tris, 1 mM EDTA, and 750 mM NaCl. Chambers were filled with the antibody solution and incubated for 30 min. Following antibody binding, the DNA-DNA closing interactions were removed by a strand displacement mechanism upon 10 min incubation with 100 µM 17 bp DNA opening DNA strands (Table S8). The closing of the nanosensor by bivalent binding of antibodies was quantified by acquiring single-molecule fluorescence scans on the confocal microscope.

### Restriction enzyme activity assay

For monitoring restriction enzyme activity, nanosensors (Version 1) were equipped with six or eight 11 bp DNA-DNA closing interactions containing a 6 bp sequence (Table S6) specific for the binding and cleavage of the restriction enzyme XhoI. For each sample a separate chamber with closed nanosensors was prepared and filled with 1×CutSmart™ buffer (New England BioLabs, USA) containing 50 mM potassium, 20 mM Tris-acetate, 10 mM magnesium acetate, and 100 µg/ml recombinant albumin. 0.5 µL of XhoI (20.000 units/mL, New England BioLabs, USA) were added and incubated for 10 min. The opening of the nanosensor by the cleavage of the DNA-DNA interactions was quantified by acquiring single-molecule fluorescence scans on the confocal microscope. For monitoring the cutting kinetics of the XhoI restriction enzyme cleavage reaction of the closing interactions, single-molecule fluorescence transients were recorded on a wide-field microscope after the addition of 0.5 µL of XhoI (20.000 units/mL, New England BioLabs, USA).

## Data analysis

A python script was used to process the acquired single photon counting data which is available on GitLab https://gitlab.lrz.de/tinnefeldlab/cospota). Briefly, the software finds single spots using a wavelet decomposition-based approach and then calculates the spotwise PR as PR = I_Red_/(I_Red_ + I_Green_). We used the spotwise PR to distinguish between open (PR<0.3) and closed (PR>0.3) conformations. Data was plotted and fitted using Matplotlib, Scipy and Numpy. For the estimation of K_½_ and n_H_, we used the modified Hill equation:

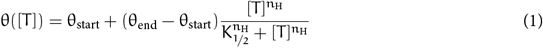

which also allowed fitting the start (θ_start_) and end (θ_end_) points of the titration curve given by the target concentration [T] and the occupancy at each concentration θ([T]). The reported errors are standard errors of the fit.

