## Supplementary Information for "Engineering Modular and Tunable Single Molecule Sensors by Decoupling Sensing from Signal Output"

†These authors contributed equally.

**The PDF file includes:**

Supplementary Text
Figures S1 to S17
Tables S1 to S9

**Other Supplementary Materials for this manuscript include the following:**

Movies S1 to S2

**Supplementary Text**

Detailed DNA origami folding procedure

16 µL of unmodified staples was mixed with 4 µL of modified staples, 5 µL of FoB20 and 25 µL of p8064 scaffold (produced in-house). Modified staples consisted of the desired modified staples and the respective unmodified staples to replace the modified staples that were not desired in the respective experiment. An exemplary recipe for the modified master mix for the 2×13 sample from Fig. 2 is given below:

Red and green dye strands (2×1 µL)
Row 4 closing staples (4×1 µL)
Row 1-3 staples without closing interactions (12×1 µL)
Biotin 1-4 (4×1 µL)
Stabilization staples without stabilizing interactions (4×1 µL)

We used two slightly different versions of the DNA origami design. Version 1 was used in the experiments described in Fig. 3 and 4 in the main text, while Version 2 was used in the experiments described in Fig. 2. The staples needed for Version 1 of the sensor are described in Tables S1, S3, S5-S7 and the staples needed for Version 2 of the sensor are described in Tables S2, S4 and S5.

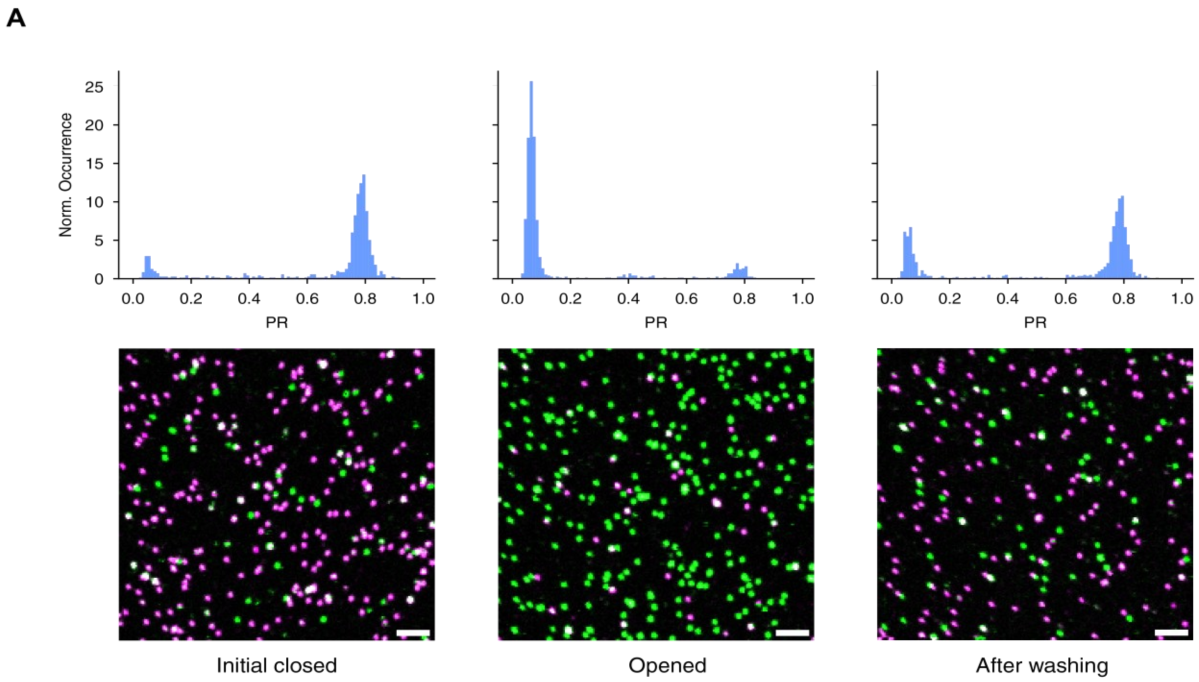


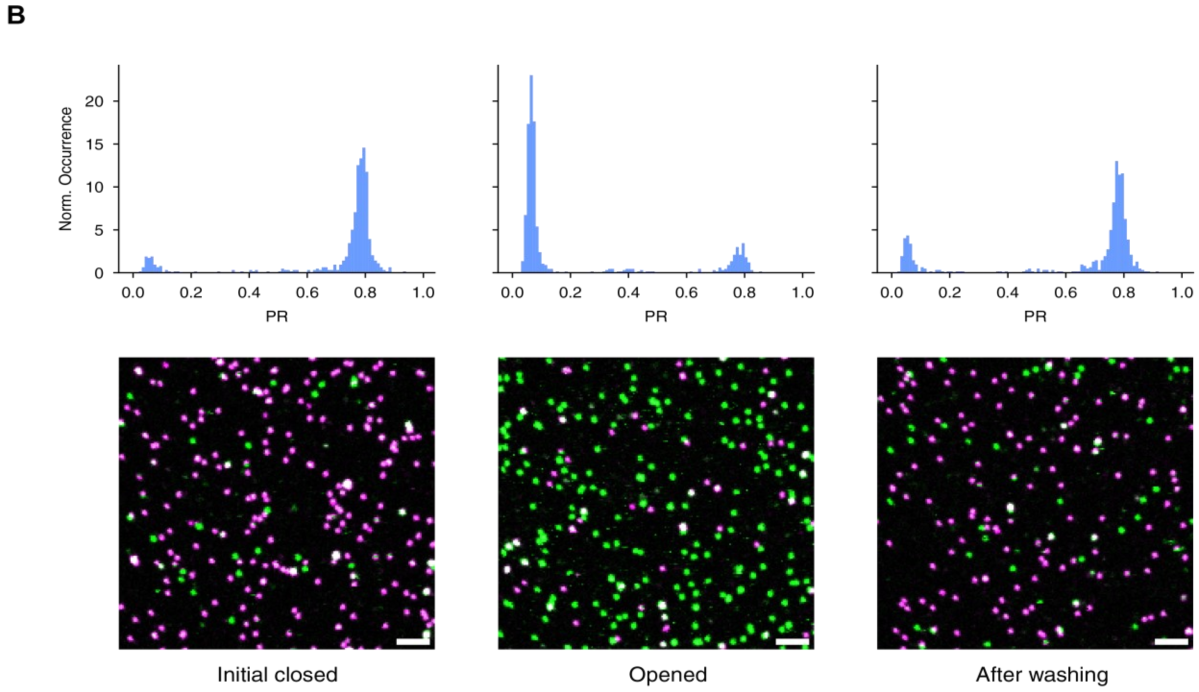


**Fig. S1.** Reclosing of the hinge nanostructure with two closing interactions measured in the presence of 50 mM (**a**) and 400 mM (**b**) NaCl. Samples were opened by incubating them with 100 µM opening strands for 1h. Afterwards, they were washed at least 10 times with the respective buffer and scans were taken after approximately 190 hours to allow for complete equilibration.

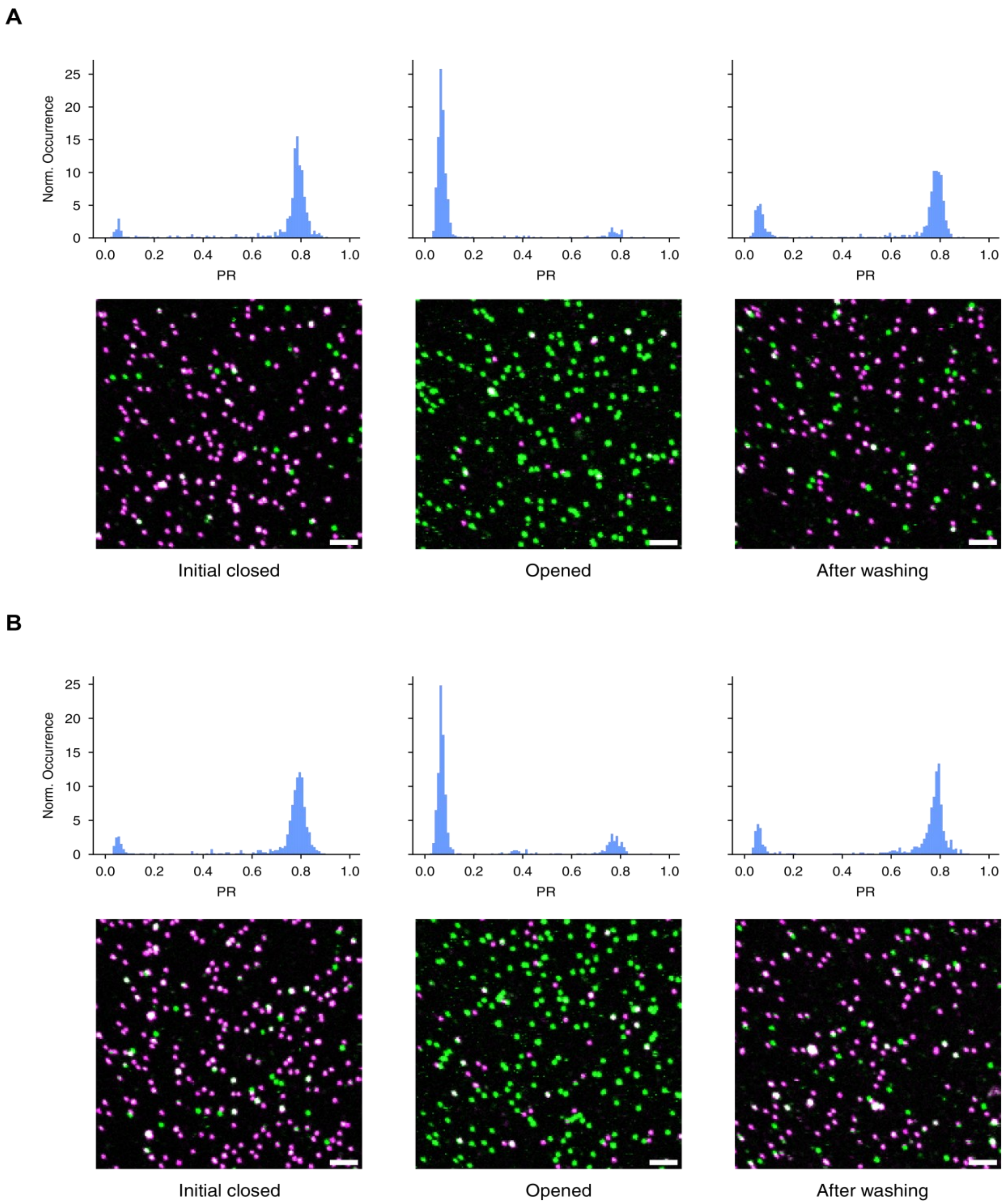


**Fig. S2.** Reclosing of the hinge nanostructure with four closing interactions measured in the presence of 50 mM (**a**) and 400 mM (**b**) NaCl. Samples were opened by incubating them with 100 μM opening strands for 1h. Afterwards, they were washed at least 10 times with the respective buffer and scans were taken after approximately 190 hours to allow for complete equilibration.

**Closed structures**

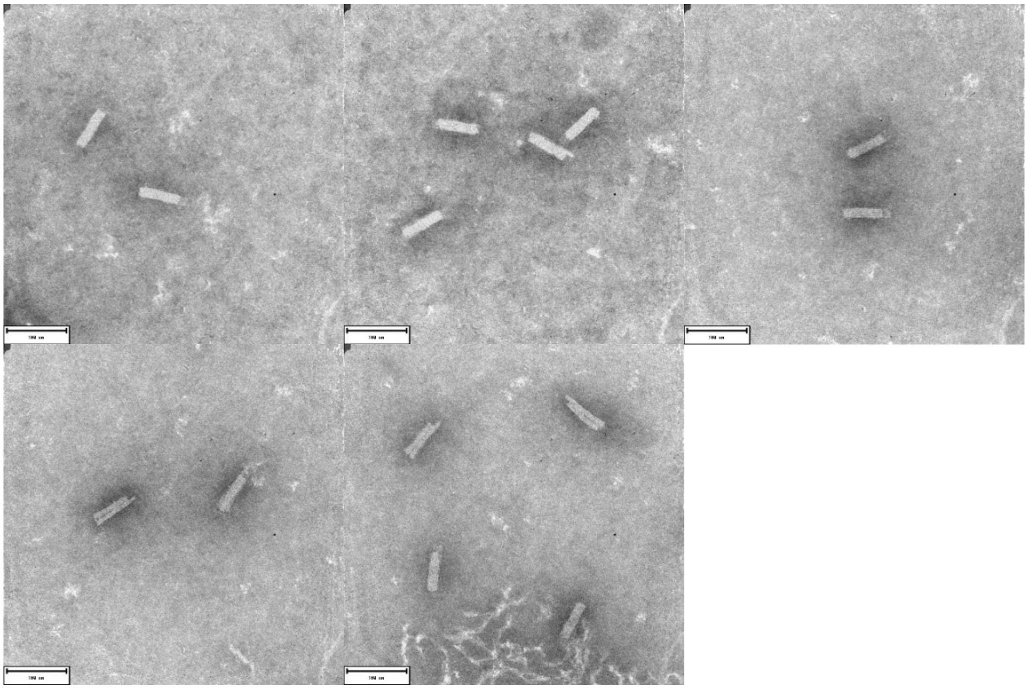

**Open structures**

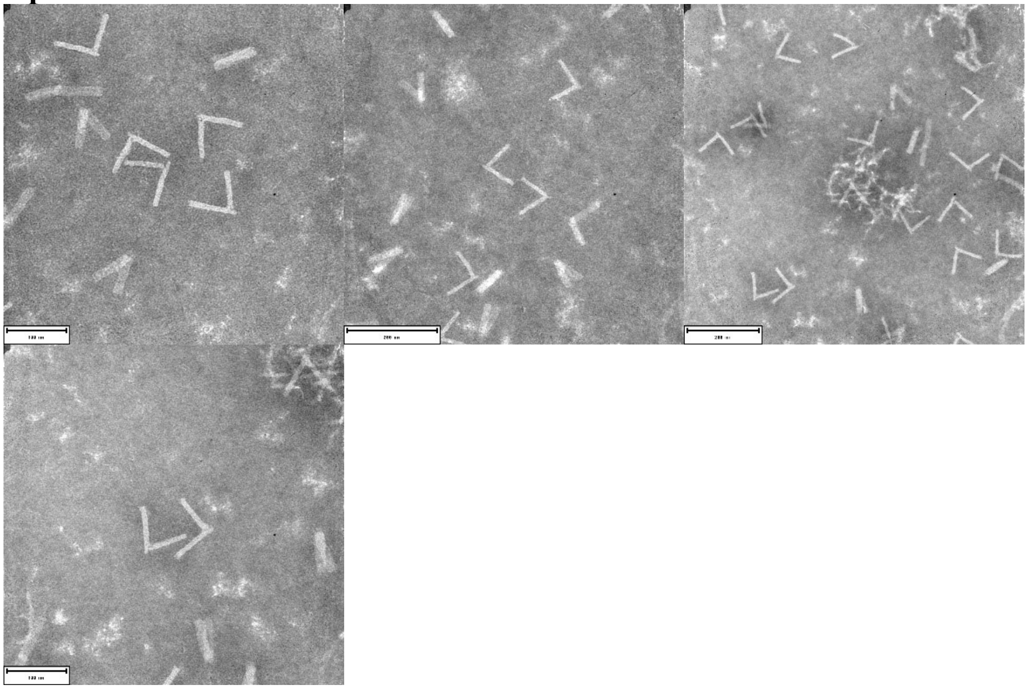

**Fig. S3.** Full field of view of the TEM micrographs used for the collage shown in Fig. 1b.

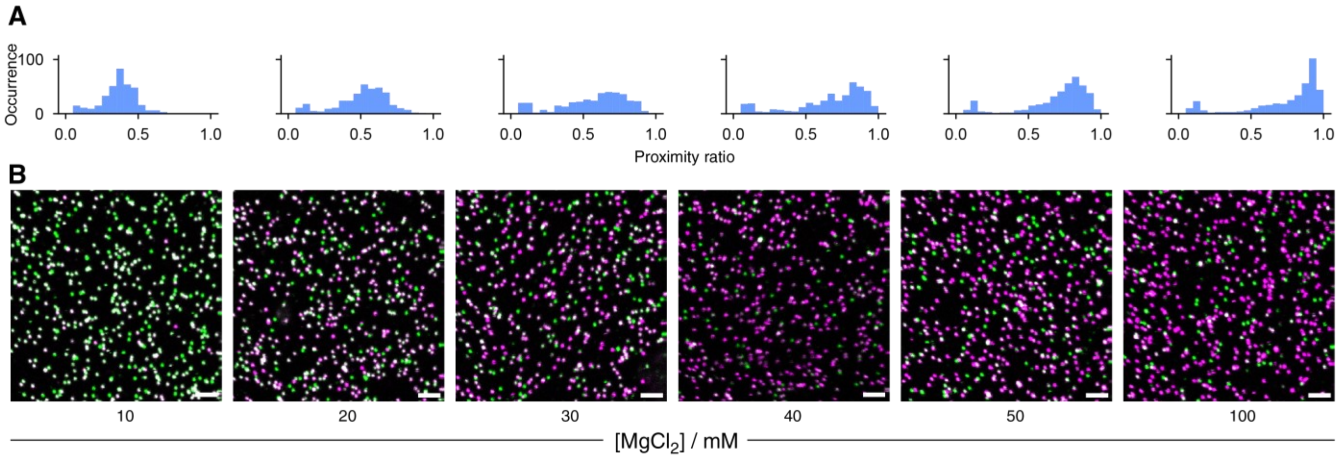


**Fig. S4.** FRET proximity ratios (**a**) and single molecule confocal scans (**b**) obtained for Version 1 of the nanosensor with a FRET pair consisting of ATTO532 and ATTO647N incorporated directly onto the hinge arms (Table S5) at different ionic strengths (increasing concentrations of $MgCl_2$). Scale bar: 2 µm.

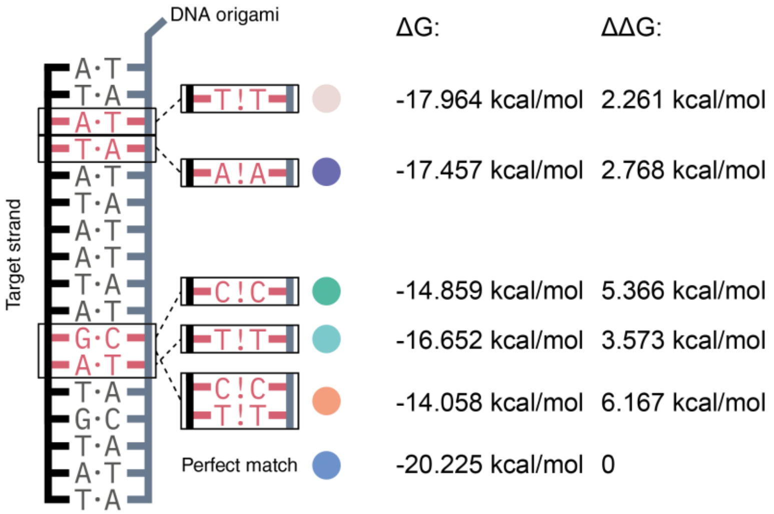


**Fig. S5.** Binding free energies (ΔG) for the different off-targets as well as for the perfect match as shown in Fig. 3 in the main text, as calculated by NUPACK at 25°C in a buffer containing 750 mM NaCl and 5 mM $MgCl_2$. ΔΔG is the difference in binding free energy when compared to the perfectly matched target.

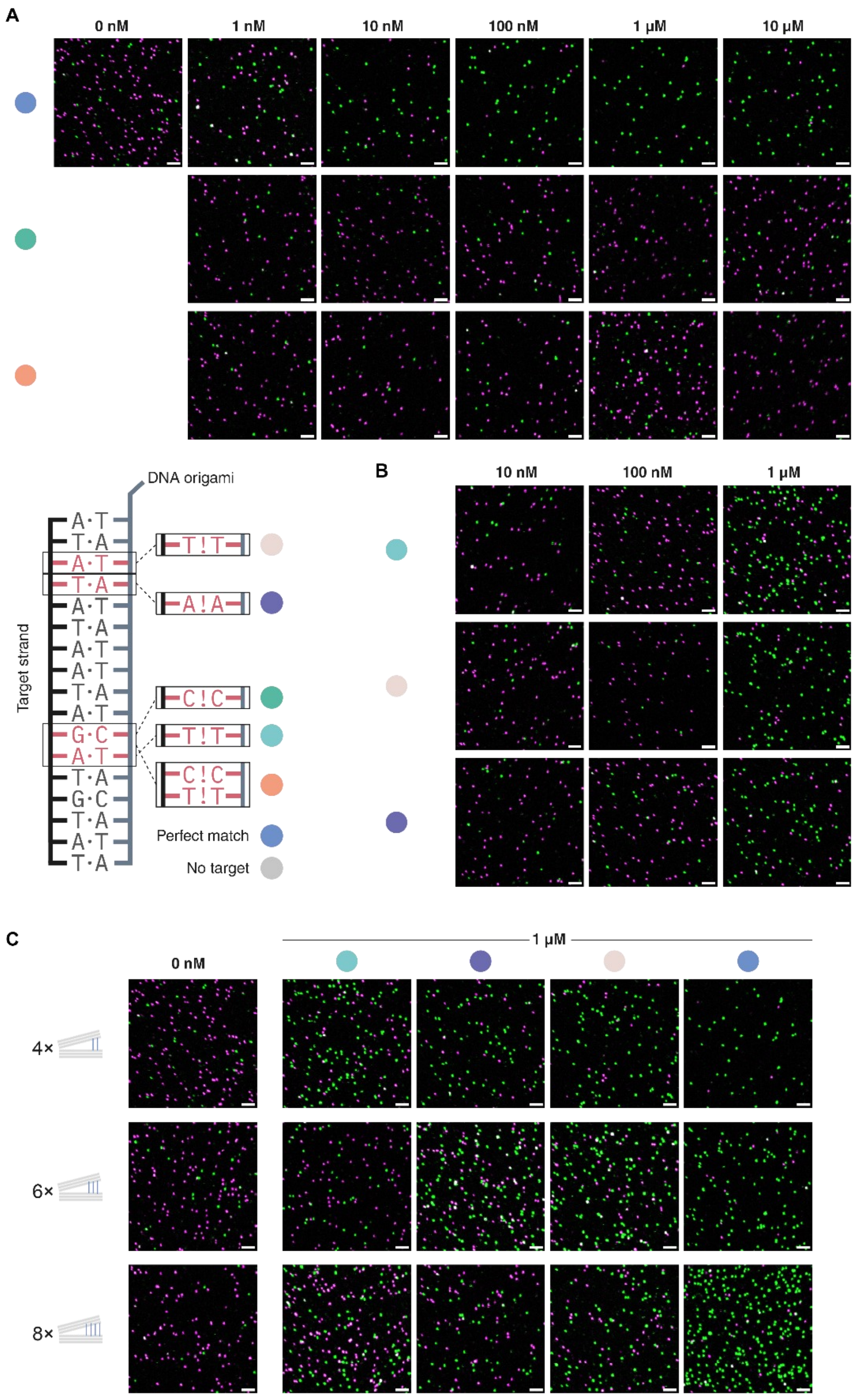


**Fig. S6.** Exemplary confocal microscopy scan images of the experiments shown in Fig. 3B (**a**), Fig. 3C (**b**) and Fig. 3D (**c**). Scale bar: 2 µm.

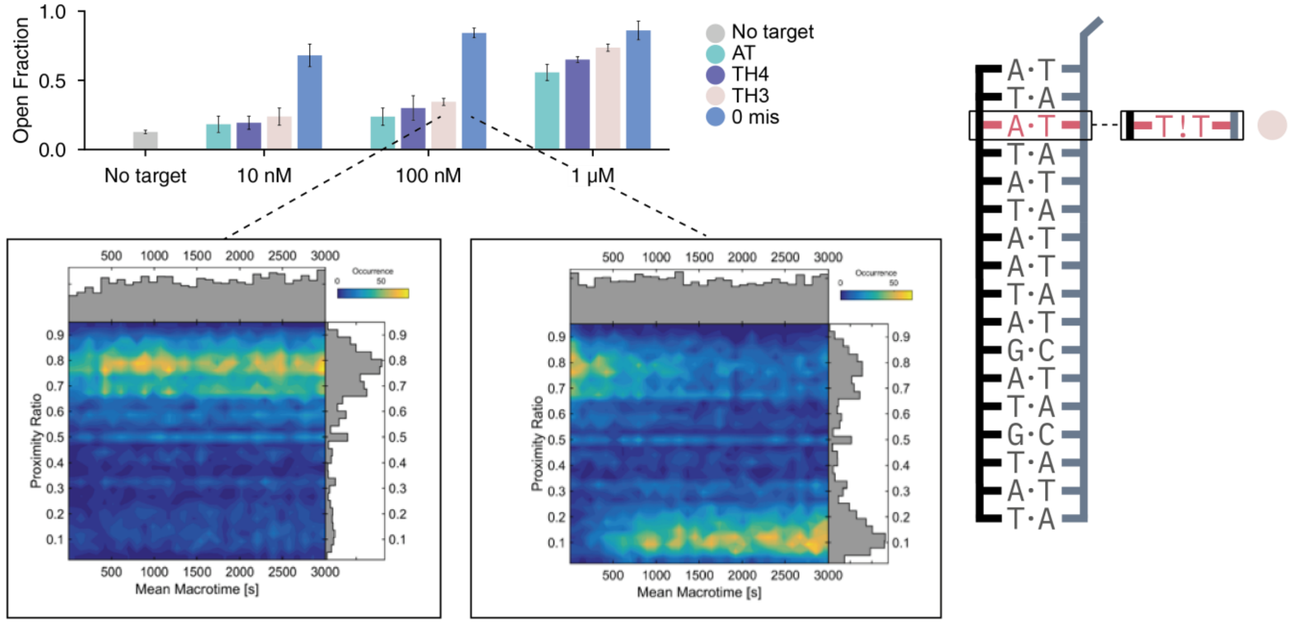


**Fig. S7.** Nanosesnor opening kinetics **(**proximity ratio vs. time) obtained for 100 nM of the target containing a single nucleotide mismatch (left) and 100 nM of perfectly matched target (right) measured at 1 nM concentration of DNA origami nanosensor containing 4 closing interactions.

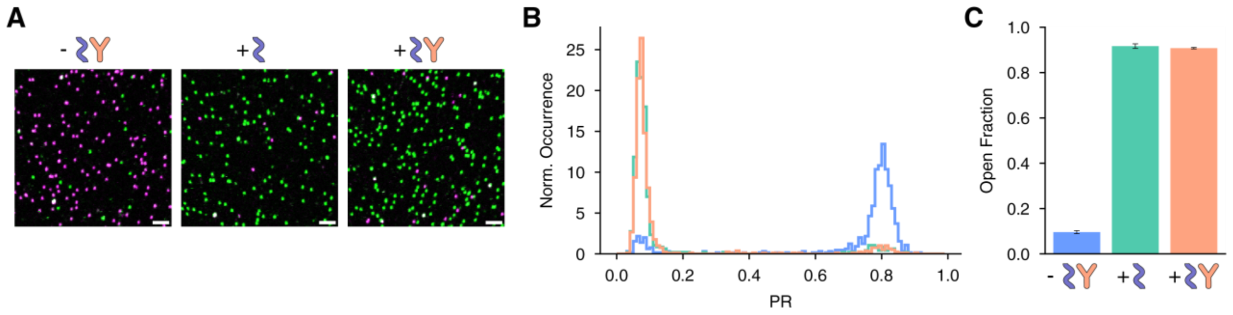


**Fig. S8.** Control experiment for nanosensor containing no Dig recognition elements. In this case, opening of the sensor is observed even after incubation (100 nM, 30 min) with anti-Dig antibodies. (**a**) Exemplary confocal microscopy scans. Scale bar: 2 µm. (**b**) Proximity ratio histograms for one experiment. (**c**) Fraction of open sensors. Error bar represents the standard deviation obtained from three independent measurements.

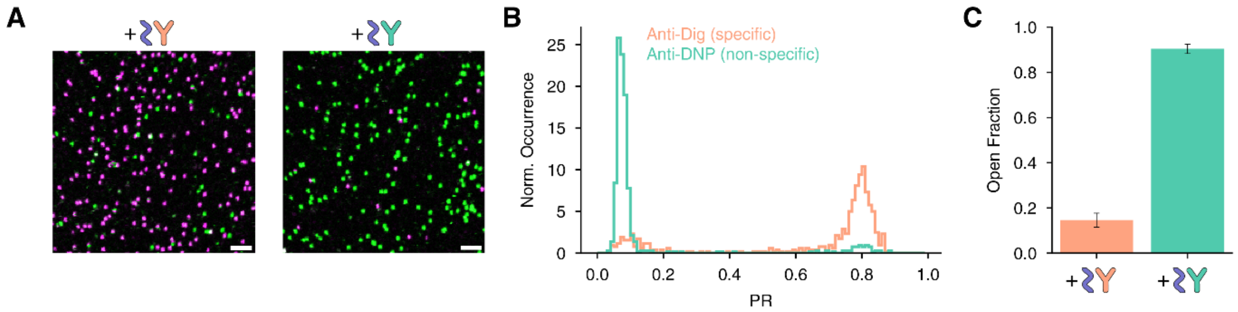


**Fig. S9.** Comparison of the extent of opening for sensor with Dig antigens when incubated with 100 nM of the corresponding specific antibody (Anti-Dig, orange) to 100 nM of a non-specific antibody (Anti-DNP, green). (**a**) Exemplary confocal microscopy scan images. Scale bar: 2 µm. (**b**) Proximity ratio histograms for one experiment. (**c**) Fraction of open sensors. Error bar represents the standard deviation obtained from three independent measurements.

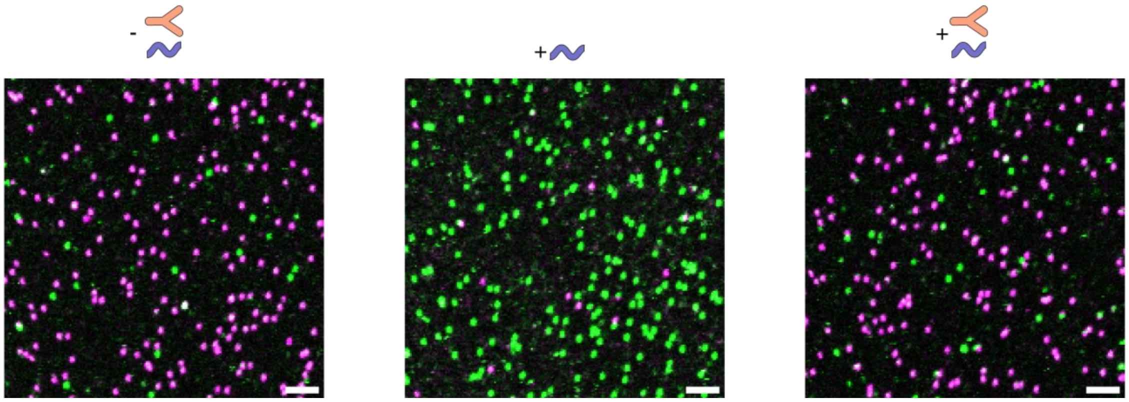


**Fig. S10.** Exemplary confocal microscopy scans for the data acquired for the antibody sensor (data shown in Fig. 4c before addition of the target (left), upon the addition of DNA opening strand alone (middle) and upon incubation of 100 nM antiDig antibody and addition of DNA opening strands (right) in heparin plasma. Scale bar: 2 µm.

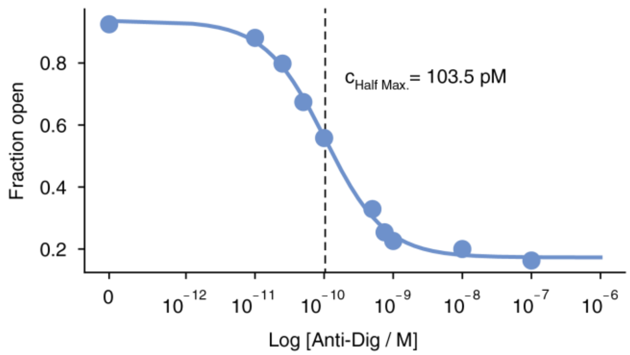


**Fig S11.** Estimation of the response window of the Anti-Dig antibody nanosensor. We achieve a $c_{Half\text{-}Max.}$ value of 103.5 pM.

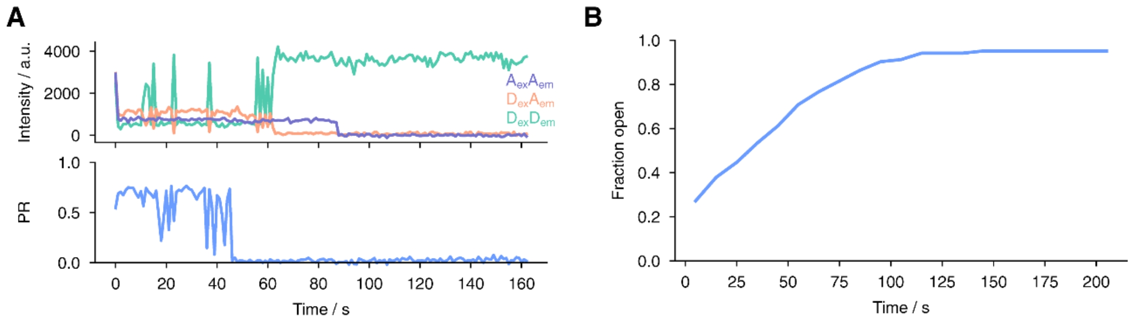


**Fig. S12.** Kinetics of the XhoI restriction enzyme reaction. **(a)** Exemplary single-molecule FRET trace and **(b)** Average of 103 single-molecule transients of the cleavage reaction of the closing interactions (66 units/mL XhoI, 10 min incubation) monitored by the DNA origami sensor. The FRET transients of the single DNA origami sensors were recorded on a wide-field microscope.

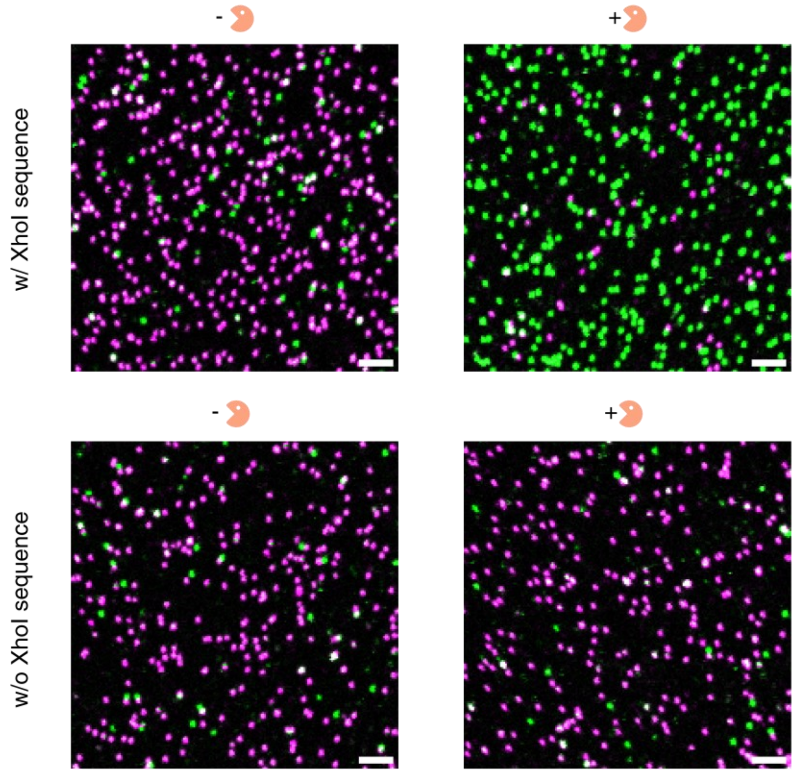


**Fig. S13.** Exemplary confocal microscopy scan images obtained for the nuclease nanosensor (shown in Fig. 4d-f). Scale bar: 2 µm.

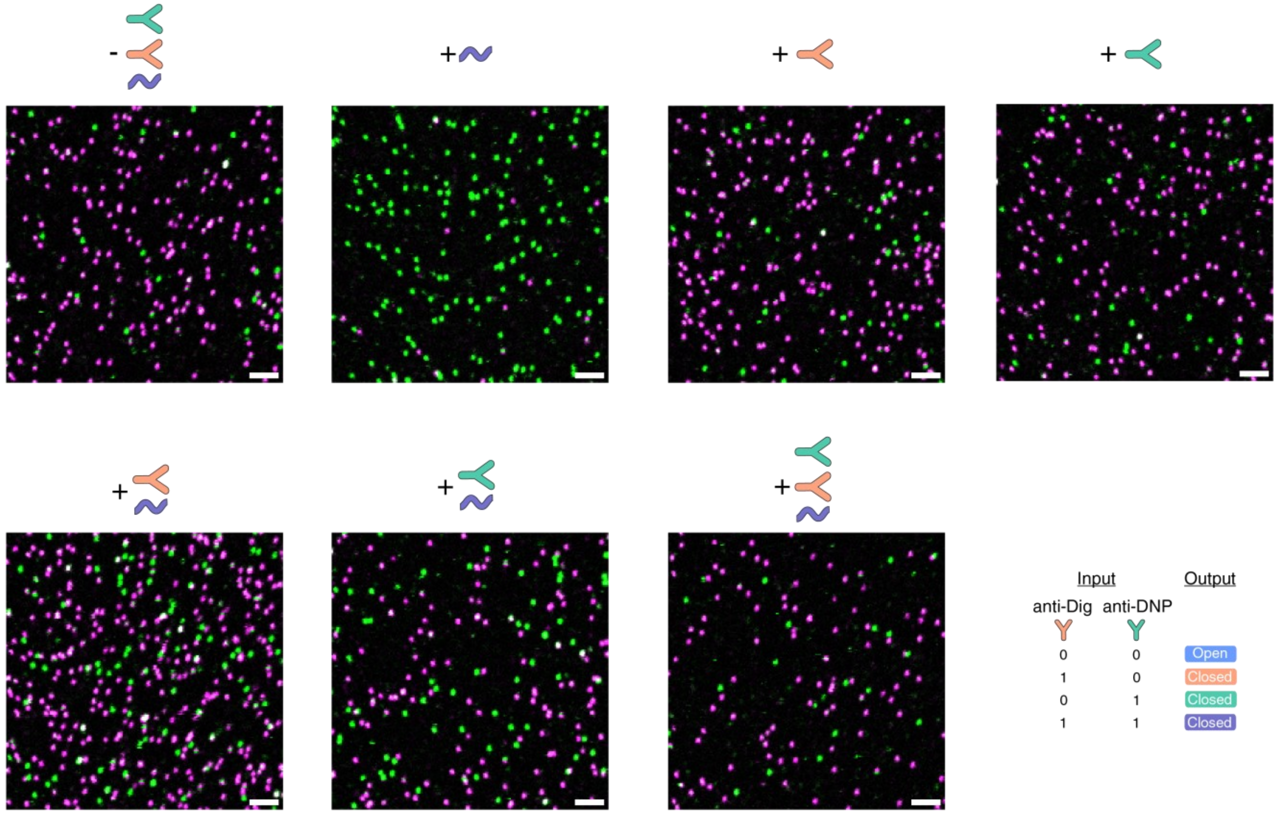


**Fig. S14.** Exemplary confocal microscopy scan images obtained for the multiplexed sensor for antiDig (orange), and anti-DNP (cyan) antibodies (OR gate) as shown in Fig. 4g-i. Scale bar: 2 µm.

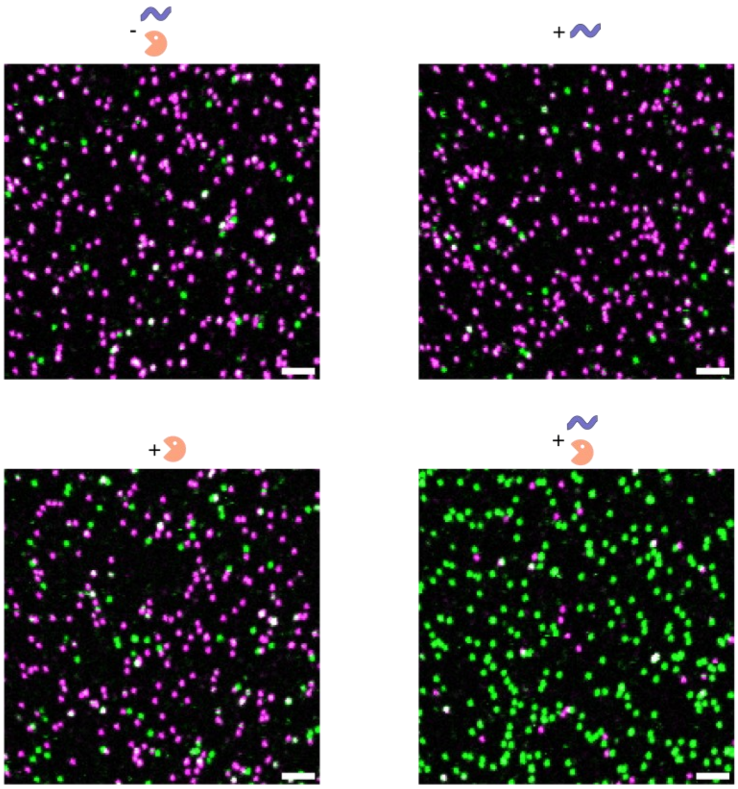


**Fig. S15.** Exemplary confocal microscopy scan images for the multiplexed nuclease and nucleic acid sensor (AND gate) shown in Fig. 4j-l. Scale bar: 2 µm.

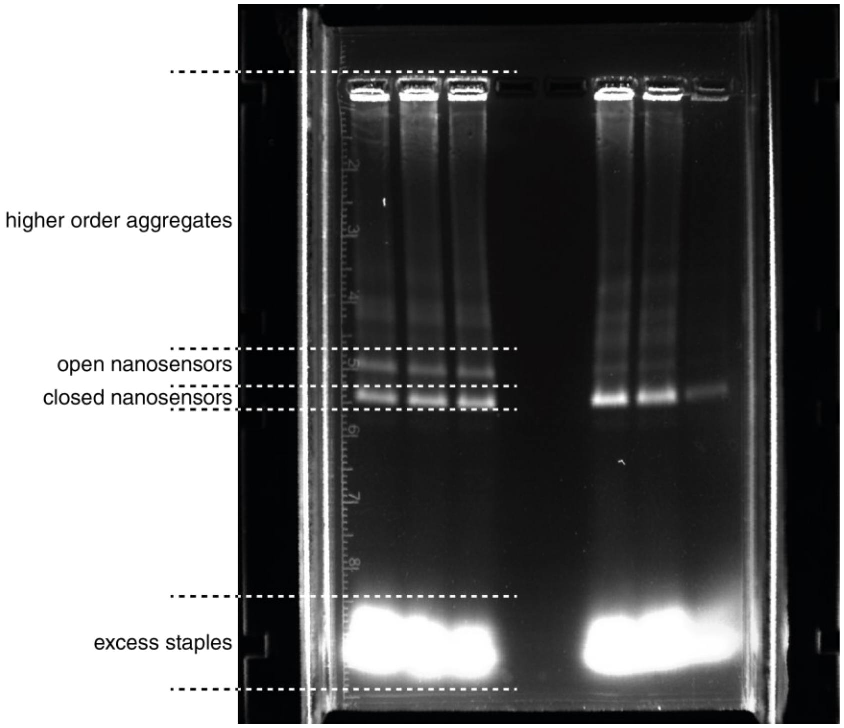


**Fig. S16.** Exemplary fluorescence scan of an agarose gel used for purification of the nanosensors. The respective band was then cut using a scalpel.

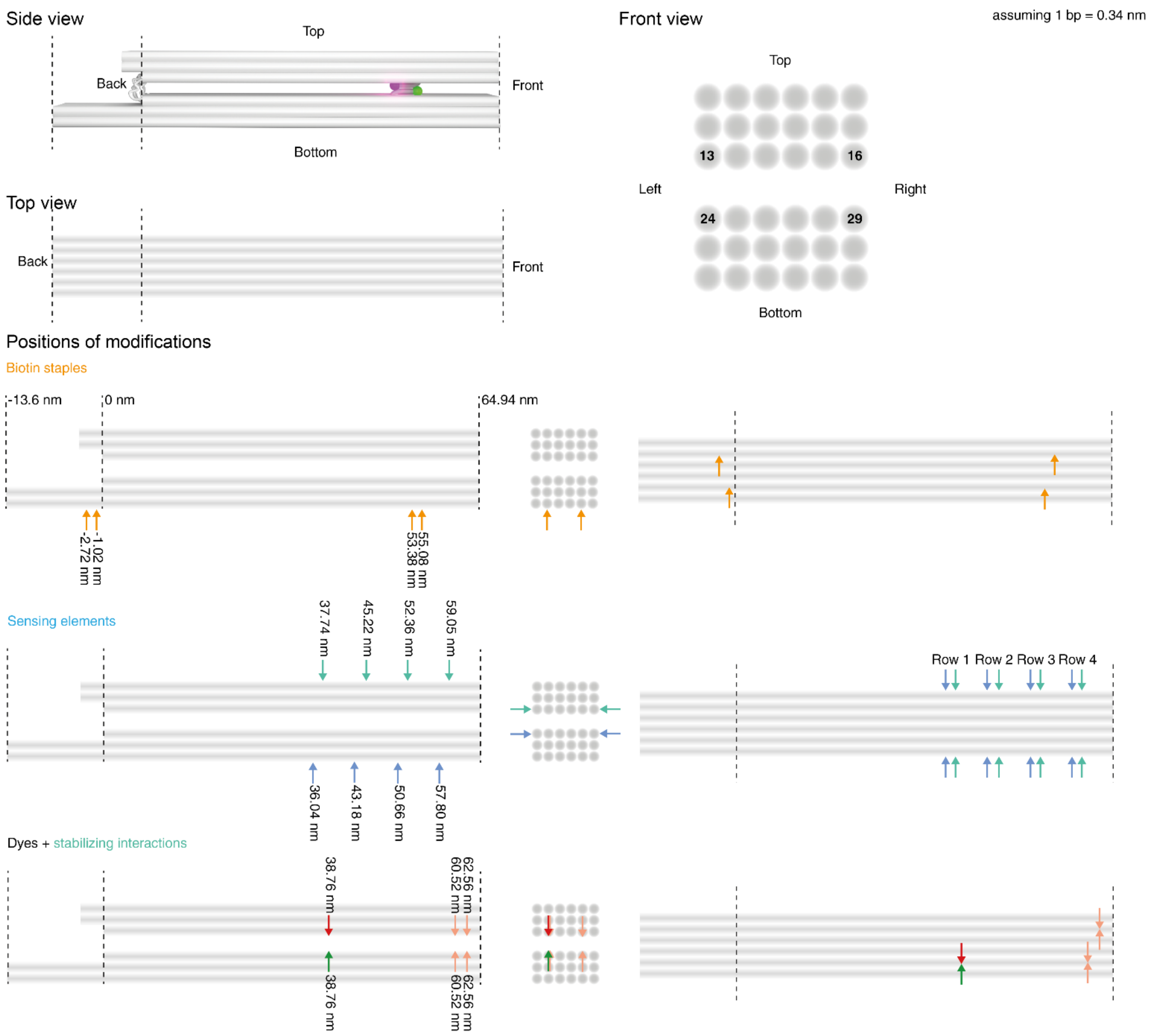


**Fig. S17.** Depiction of the placement of the modifications (FRET pair, sensing elements, biotin staples used for immobilization as well as stabilizing interactions) in the nanostructure. The numbering in the front view represents the helix numbering in caDNAno and are also used to indicate the positions of the sensing elements in Tables S3, S4, S6, and S7.

**Table S1**. Unmodified DNA origami staple sequences used for Version 1 of the nanosensor.

| ID | Sequence (5'->3') |
|---|---|
| 1 | AAATATCCCATATGTTCAGGCAAATCCAGTGATGACAGTTGGGCGGTTGTAAATC |
| 2 | ATATTACCAAATACCTGTAATTCTAAAACTTT |
| 3 | GCTGGGAATCAAAGAAACGTAGACGGGAGAATTAACTGAACACCCA |
| 4 | GTCCACTAAATCCCTTTAAACCAAGGCGGGCCTGT |
| 5 | ACGTGGACTCCAACGT |
| 6 | CAAAGGGCTCAAAAATCAAGCCGTGTATGTTAGCAAAGGTGCTGCGGC |
| 7 | GAGCACGTATCCAAATTTACCGCGAACTGGCAAAGAAACGCAAAGACATTAC |
| 8 | CGTGGAACTCAAACTATCGGAAATGGATAAGAG |
| 9 | AAATTAGTAATAACATCATCACACGAATTTAGGCACCG |
| 10 | TTTACGAGCGTGGCGTTTTATAGCCGATTTACCAGCGCCAAAGTATC |
| 11 | AGGGCAAATTAACCGTTGGCCAACAGAGAATCGCAGAA |
| 12 | CAGCCGAAATCGGCAATTAAAGA |
| 13 | CACCCAGCAGTGTTTTGAGGCCACCGAGTAAAGCGTAAGAAAGCCAAAAGCCTG |
| 14 | TATATAAATCAAAAGAAT |
| 15 | CGCTAAACAACCCTAATTGGAAAAACGCTCATGGGCCAGCCAAGAACAAGGAAAATAG |
| 16 | TATAAACAGCCAAAGTCA |
| 17 | CTAACTTCTTTGCAGGAGGCCGATTAAAGGGA |
| 18 | ACTCCTTAAAGCGCATATTTTTGTTTAACGGAAAAACCCAGAACA |
| 19 | AAAGCCCAATAGCCAGTAATTATTTACAACAGAGGTGAGGCGGTGCTG |
| 20 | GCTTTTGTCACAATCAATATTAG |
| 21 | CCAGCAGGCGAAAATCCTGTTTGATGGTGGTCAGAATGCGTAC |
| 22 | TCCTTTACAGAGAGAATAACATAAAAACAGGGTTACGCATTTTATTTAGGT |
| 23 | AACATATTATTTATCCCAATAA |
| 24 | CGCACAGGGTGCACTCTGTCGTAGAAAATACATACATAAAGAAA |
| 25 | CCATCGCCGTAAAAAAAGC |
| 26 | CATTGACCTAAAAATTACCTTCAAATATAATTTTAAAAGTTTGAGTAACA |
| 27 | AGCTGTGATAACAAAATTAAGTTGGCACTCGTATT |
| 28 | ACCCACCGGAAAAAGAAGGGAAGGTTGAAGTATTAGACTTTACAAACAAAACA |
| 29 | AGACGATCCAGCGCAGGTTTTCACGGTCATAC |
| 30 | CACTGCGCGCCTCGGCCTTTAATCGCAGTAAA |
| 31 | GCAACCGACAAATCATCGTAGTAATATCGTCTATCACTTTGAC |
| 32 | GAATTTCGAGCAAGCAAAAGTAGAACTTTCCTC |
| 33 | CATTCCCTTTTTAAGAAAAGTTAAGCCCATCTTACCAACGCTATAGAC |
| 34 | CTCTCTGAATAATGGAAGGGAGCGGAGAATTT |
| 35 | ATATTCACAAACAAATGTACATCGTTGCTTCTAGACAAAGCATAG |
| 36 | GCTCATTTGCCGCCAGAGGGTAAAGTTAAACG |
| 37 | CACCAGACGATATCAAAATTATTTGCAATCATTTTCTAAAGCA |

| | |
|---|---|
| 38 | GTCTCATTTGTTTAATGGAGATAAATTGGCAGATTCACCAGCTTGCCTGTCAGA |
| 39 | TATCACCAGTAGCACCACCACG |
| 40 | TAATTTCATTAAAGGTGAATACAAAAGGAGTAGGGCTCCCGACTTCCATCAC |
| 41 | CCGGAACACCGGAACGCTTTGAA |
| 42 | TAGTGAACAAGAAAAATAACCAATCAAACGGG |
| 43 | GTCTGAGAGACTACCTTTTAGA |
| 44 | AGATTTGAAATAGAGGCATTAAGTTTAAATAATAACGGAATACCCTG |
| 45 | ATCGGTTAAATACATATTTATCATATGGACAAAGTTACC |
| 46 | CCTAGGAATTGAATGATGAACTTTTACAGCCACCCTCAGAGCCACC |
| 47 | CGCCCCTTCTGACCTGAAAAGAGTCTGTGCGGGAGTATCCTGAAATAATA |
| 48 | CGCCTGGCCCTGAGACGCGTGCCATAGCGTCTGCTGTAGTCGAGCTTGATAAATT |
| 49 | GGGTGGAATTCGTAGCGAGAGATAACAGTCAGCAGCGTAAAGGAATTGCGAATTTTT |
| 50 | GGGGTTTCTGCCAGCAGAGTTGCAGAGTAGTA |
| 51 | TTCCAGATGCCGGGTTACGCAAACGCAACGCAAGAAAACAGGGAGAGATCTTAGGATT |
| 52 | CCGAGCTCGTTTTTCGATTTTAAGAACTGGCAGGGAACCCATGAGGA |
| 53 | TGAAACTCGTTTATAGATTTAGTTAA |
| 54 | CGGATCCGCCGGGCGCGGTATGCCAACAAATCATACGTC |
| 55 | TGGGGAATGCAGCATTTGGGACAA |
| 56 | GCATGCGTCCGTGAGCCTCCGCTGATTGTCAACTT |
| 57 | AAACCATTCCATGCTTTTGGACTTTTTGAACTGACCAACTTTGAA |
| 58 | GGTACGAACGAGCCAGACGAGGTAAAATACGAGGCGCAGACGGT |
| 59 | GCTGATTGCCGTTCCGCTGCAGCCGAATATAACAATA |
| 60 | TCGTCGCTAAATCGTTTATGCAACTAATAGTAGCCCGAAACTACAGAGGC |
| 61 | GCCTGTAATACTATTGTAAATATGGGATCGGATAAGTGCCGTC |
| 62 | ACGTAAATCGGTCATTAAATTCTTTCCAGTATAGCCCGGAATAG |
| 63 | GGAAAAAATTCGTGTACCAGGAACAACAAAGACAGCATCGGAGGAC |
| 64 | TTACCAGTATTCGCCAGACGACGACAGTATCTGATATAAGACGTTA |
| 65 | GCGAATCATTTTTAATAAAGCTTTTCACGCCGCTTTT |
| 66 | AAAACGAGCGATTAATCTCCGTGGGAACAAAGAGCCACTGA |
| 67 | CTCTGAATTTACCGTTACGCAGAAAAGCCCCAGATAAAAATTCTA |
| 68 | GTAAGGTACTGGTAATAAGTTGAGAAGGATACAAAGG |
| 69 | ATGGCTTTCCGGCACCTGGTGAAGATATTTAATTTGCGGGTTCAA |
| 70 | CTGGCACTCCAGCCAGCTTTTGATGATAATAAGCAAGGATAGCTTTTTCGTC |
| 71 | TCTGTAGCAATAGAACGCGACTTTCCAGGAGAGATAACCCACAAGA |
| 72 | GTGTATTTTTGAATGGCTAGGAGCACTTATTCATCCTGATTCGCCTCCC |
| 73 | CCGTGCAACCGTACTGCGTAACGATCTAAAGAAT |
| 74 | CGTCGGATGTTGGGTAACGCCA |
| 75 | CCCTGAGAGTATTAAGAG |
| 76 | TTAATGAAGATCGTGCCGGAAACCAGGCAAAGCGCCCCC |
| 77 | CAATCATAGTTACAGGAGGTTTAGTACCGCCACCCTTTGGTGTCATC |
| 78 | AAAAACGGATAAGCAGCAACCGCAAGATGCGGTATCATTTCGCGTAA |

| | |
|---|---|
| 79 | TGGCACAACGCCCCACCCTCAGAACCGCCACCCTCACGGCGGATAGC |
| 80 | CAGCGCCAAGCTCCGTCGGTGGTGCCATGGCGCTTTTATATTTTATACATAATAAA |
| 81 | TAGCATGTCAATCATAAAATGCAAGTCAAATCCAGGTC |
| 82 | AGAATTAGATAGAGCCGGCGCTCACAATTCCACAAGCTGCATACGAACTA |
| 83 | TCTACTAACATTAAAGTAACACCACCCTCATTTTCAGGGATAG |
| 84 | AACATTTATTTCGGTCCGTTCTCACGGAAAAAGAGCCA |
| 85 | CCCATCCAATGGCAGCATTCAGAGGTGGAGCCGGCCAG |
| 86 | AGCCGCACTCAAAGCATAAAGTGTAAAGCCCG |
| 87 | AAGAGGTCATTTTTGCGAATCCCCGGTCTTTA |
| 88 | CTGAAAGCGGAGACCTTCATCAAGAGTAATCTTGAGAG |
| 89 | GGGTAAAGTACAGCAACGGGACTTCAA |
| 90 | AACCTTGTTATCGTCACTGTTGCCCTGCTGGTGCTGAATTAAGCTAAC |
| 91 | GAGCAACCTAAATCCGCGACCTGCTCCATGTTACTAATCTACGTCAT |
| 92 | ATTCAACTTGCCTAATAGCGGGGT |
| 93 | AAACATCATAAATATTCATTGGATGGCTACTC |
| 94 | AATTGGGCGGCTGGCTTTGCATCAAAAAGATTTTTTAATCTC |
| 95 | TAATCATTGACAGATGAACGGTGTA |
| 96 | ACGGAACATGTCGAAAACGAAAGAGGCAAAACGCATAACAATAA |
| 97 | AAGGCTTGCCCTGACCAAGAACCCCTGACT |
| 98 | CAGACCAGGCGCATATTGAGATGAGACTGGTGTTC |
| 99 | AGAGGTGAATTAGTTTTGCCGAGGATCCACTGGTGTGTTCAGCGGCA |
| 100 | CAATCATATCATTATAACCAAAATAATCATGGAGGTTTCTTTGCTCGTCGGA |
| 101 | GAAACAAAGTACAGAGATTTA |
| 102 | CGGATATTCATTACCCAAATCAACTGCTCATCTTTAAAC |
| 103 | TTTGAACGAGGGTGGTGTCTATGACCCTCCGGCCAAGATAGACTTTCTCCGGCTT |
| 104 | TAAACAAAAGAACCTTATGCTTTTCACCAGTGAGACGGGCAACATCACAGTTAGAGG |
| 105 | AGTTTCCAGTCACCCTTGATTCCCATAAAGCTGCCGGACTCGCCATGT |
| 106 | CCACTACGTGCAGGGAGTTTGACCATTAGCAAGTCTG |
| 107 | ACACTTTGTATCATCGCCTGATAAATTGACATTATGGAATTACGAGC |
| 108 | GACCCCCATTGCGCCGGCGCGAGCGCATCAAT |
| 109 | ATAAATCAGTACCTTTTTTGATTAATGTGT |
| 110 | ATATCGCGAAGAGGAAAAATGTTTGTTTAATTCCCTTCAC |
| 111 | GCGGGATCTTAAACGCGATAAAACCAGTCAGTTTGCGTATTGGGCGCCA |
| 112 | AGGTTGAAAATCTCCAAAAACAGCCCTAGGAACGCAGATGGGCCGGTGCG |
| 113 | TGATTTCGAGGTGAATTTCCATGTACCTGTGAGCG |
| 114 | AGGATTAGAGAAAAATCACTCAAATGTCAGTGAAT |
| 115 | CAAACCATCAATATGATATTC |
| 116 | GCTCAAAGCGAACCAGACCCAGAAGCCGGAATCGCCAGAACGCAAGCGG |
| 117 | AATGCCGGAGAGGACTAGAAGCCTGTTTAAAAACG |
| 118 | CAGTTTCAGCGGAGTGAGATTTTCTGCG |
| 119 | AAGGTCGGTCGCTGAGGCTAAGGCACCAACACTATTAATAAATAATGAA |

| | |
|---|---|
| 120 | GAAAGGCAATCGATGGTAAAACACAGTTAA |
| 121 | TTCGTAAAAGGAAACAGTAGCGACCGGAAACGTCACCAATGGTG |
| 122 | TTAATCTAAAATATCTTTATTAGTCTTACTAGAAACGCTCAACGCGA |
| 123 | CTATCAGGTCAACCGTTTTTAGAAATTGCTAGCGGTGC |
| 124 | GTAAATGAAATAGAAAAAAACATTGGAAGTTTATCCCTTACCCGGGTA |
| 125 | GTCGTTTTGTTATTTGAGGGTTCAGGCTGCGCAACTG |
| 126 | GCATTCCACAGAAAAGGCTGCAA |
| 127 | GTTTCGTCACCAGCGGTTTATAGCATTAATGTTT |
| 128 | GCTGAGACTCCTCAATTAACGGGATCAGAAACAGCGGA |
| 129 | AGCGGGGTTTTGCTCAGTAC |
| 130 | CAGGTTTGCTAAACAGTAGCTATTTTTAAGATTGTCAGGAGTCGTCATAC |
| 131 | GAGAGGGTGGCCTCAGATTTTGTTTTTGTGAGGAGCACATCCTCATAACAT |
| 132 | GTGTATCTCTGCCAGAATCAGCACGTACAGTGTAGAACGTCAGCGGGCT |
| 133 | CTATTATTCTGAAACCCGTATAA |
| 134 | AGTCTGGAGCAAACAAGAGCGGAGACATGCCTGAG |
| 135 | CGCTGTAACCTCCATGCTGATCTAATGCAGAAC |
| 136 | AGAGTTCGCTATTATTATACTATTAAAGCCAGAATGGAAAGCGCA |
| 137 | ATCAAAGCCAGCAGCAAAATGTGAGTCCTACCATTGGCCTTG |
| 138 | TCACCTTCAGTATTATTAATTTATAAAGTACATATAATGATTAAG |
| 139 | AACCTTTTTAATCAGAAATAATTGACAGGAGGTTGAGGCAGGTCGTAATCAGTAC |
| 140 | TCATTACATTTTAACGTCCACCAGAA |
| 141 | TCGCAATAGTATTATCATCATATTCCTGATTATCTTG |
| 142 | ATAAATATTTTAGACACCGCCACGCTCAATCGTCTGCCTT |
| 143 | AATCCTGAAGATGATTTAGATTAAGA |
| 144 | TTTGGAATTAATTTTCCCTTATATGTAAGGCT |
| 145 | AGAAGAATAACCACATAAAAAAATCATAGCAG |
| 146 | GCGTAGATCGTTATTCAAACCCTCAATCAATCCAGC |
| 147 | ATACAGTTTCGACAAAATCAACAGTTGAAAAAAC |
| 148 | GTACACAAACATGTCATAGCCTTGAGCCATTTGGGAAGAAAATACAACGCCATT |
| 149 | CGGATTCGTTCAATTATTCA |
| 150 | GCGTTCCAAGATAATCGGCCAGTTTGGAACAAGA |
| 151 | GACGACAAACTCATCGTTGCAACATACGAGCATGTAG |
| 152 | AAAACATTTTGTGCAACAGTGCCACGCTGAGAATAACGCGAGGTCCAGAC |
| 153 | AATCATCTTCTAGCAAGGCAGAATCAAGTTTGCCTCCGCCAGCAAGAAATT |
| 154 | TGTACCAGTAATAATACCGAACGAACCAATCTGG |
| 155 | TTCAAATATCAATATTGAAAAATGCGGAACAAAGAAACCACCAGAAGGTT |
| 156 | ACCGCAGCAAAAGCGCGTTTTCATCGGCAGAGCCGCAGATGAAT |
| 157 | TAAAGTCACCGACCCCTTATTAGCGTTTCAGA |
| 158 | TTTAGTATGAGGCGAATAACAACTAGCCGTCAATAGATAATACATTTG |

**Table S2.** Unmodified DNA origami staple sequences used for Version 2 of the nanosensor.

| ID | Sequence (5'->3') |
|---|---|
| 1 | GCTGGGAATCAAAGAAACGTAGACGGGAGAATTAACTGAACACCCA |
| 2 | TCATTACATTTTAACGTCCACCAGAA |
| 3 | CAGCCGAAATCGGCAATTAAAGA |
| 4 | CAGGTTTGCTAAACAGTAGCTATTTTTAAGATTGTCAGGAGTCGTCATAC |
| 5 | CTCTGAATTTACCGTTACGCAGAAAAGCCCCAGATAAAAATTCTA |
| 6 | GTACACAAACATGTCATAGCCTTGAGCCATTTGGGAAGAAAATACAACGCCATT |
| 7 | CAACATACGAGGCATAAAATGGTCCGATATATAGCCTTTAATTGTATTA |
| 8 | AAGGCTTGCCCTGACCAAGAACCCCTGACT |
| 9 | AACATTTATTTCGGTCCGTTCTCACGGAAAAAGAGCCA |
| 10 | TTTAGTATGAGGCGAATAACAACTAGCCGTCAATAGATAATACATTTG |
| 11 | CATTCCCTTTTTAAGAAAAGTAAGCAGAGCGAACCTTAATTGAGATAGAAGAACTGATAGC |
| 12 | CGCACAGGGTGCACTCTGTCGTAGAAAATACATACATAAAGAAA |
| 13 | AATGGGATAGGTCACGCAGAACCGTGTA |
| 14 | GCGTAGATCGTTATTCAAACCCTCAATCAATCCAGC |
| 15 | GCGGGATCTTAAACGCGATAAAACCAGTCA |
| 16 | TTACCAGTATTCGCCAGACGACGACAGTATCTGATATAAGACGTTA |
| 17 | GAATTTCGAGCAAGCAAAAGTAGAACTTTCCTC |
| 18 | GGTACGAACGAGCCAGACGAGGTAAAATACGAGGCGCAGACGGT |
| 19 | ATAAATCAGTACCTTTTTTGATTAATGTGT |
| 20 | CACTGCGCGCCTCGGCCTTTAATCGCAGTAAA |
| 21 | AGTTTCCAGTCACCCTTGATTCCCATAAAGCTGCCGGACTCGCCATGT |
| 22 | GCATGCGTCCGTGAGCCTCCGCTGATTGTCAACTT |
| 23 | GCATTCCACAGAAAAGGCTGCAA |
| 24 | GAGAGGGTGGCCTCAGATTTTGTTTTTGTGAGGAGCACATCCTCATAACAT |
| 25 | CGTGGAACTCAAACTATCGGAAATGGATAAGAG |
| 26 | CCACTACGTGCAGGGAGTTTGACCATTAGCAAGTCTG |
| 27 | AAACATCATAAATATTCATTGGATGGCTACTC |
| 28 | AACATATTATTTATCCCAATAA |
| 29 | ATCGGTTAAATACATATTTATCATATGGACAAAGTTACC |
| 30 | GTCTGAGAGACTACCTTTTAGA |
| 31 | CCGTGCAACCGTACTGCGTAACGATCTAAAGAAT |
| 32 | CCCATCCAATGGCAGCATTCAGAGGTGGAGCCGGCCAG |
| 33 | TGGCACAACGCCCCACCCTCAGAACCGCCACCCTCACGGCGGATAGC |
| 34 | GTCGTTTTGTTATTTGAGGGTTCAGGCTGCGCAACTG |
| 35 | GCTTTTGTCACAATCAATATTAG |
| 36 | AACCTTGTTATCGTCACTGTTGCCCTGCTGGTGCTGAATTAAGCTAAC |
| 37 | AACCTTTTTAATCAGAAATAATTGACAGGAGGTTGAGGCAGGTCGTAATCAGTAC |
| 38 | AATGCCGGAGAGGACTAGAAGCCTGTTTTAAAAACG |

| | |
|---|---|
| 39 | AGGGCAAATTAACCGTTGGCCAACAGAGAATCGCAGAA |
| 40 | AGCCTAATTTGCCAGTGAGCTA |
| 41 | AAACCATTCCATGCTTTTGGACTTTTTGAACTGACCAACTTTGAA |
| 42 | CGGATATTCATTACCCAAATCAACTGCTCATCTTTAAAC |
| 43 | ATCAAAGCCAGCAGCAAAATGTGAGTCCTACCATTGGCCTTG |
| 44 | TCGTCGCTAAATCGTTTATGCAACTAATAGTAGCCCGAAACTACAGAGGC |
| 45 | GAAAGATTCATCAGTTACGGAGATCATCTTT |
| 46 | ACACTTTGTATCATCGCCTGATAAATTGACATTATGGAATTACGAGC |
| 47 | GCCTGTAATACTATTGTAAATATGGGATCGGATAAGTGCCGTC |
| 48 | CAATCATAGTTACAGGAGGTTTAGTACCGCCACCCTTTGGTGTCATC |
| 49 | CTCTCTGAATAATGGAAGGGAGCGGAGAATTT |
| 50 | GGGTAAAGTACAGCAACGGGACTTCAA |
| 51 | AAGAGGTCATTTTTGCGAATCCCCGGTCTTTA |
| 52 | TAGCATGTCAATCATAAAATGCAAGTCAAATCCAGGTC |
| 53 | TTTGAACGAGGGTGGTGTCTATGACCCTCCGGCCAAGATAGACTTTCTCCGGCTT |
| 54 | ACGGAACATGTCGAAAACGAAAGAGGCAAAACGCATAACAATAA |
| 55 | AGCCGCACTCAAAGCATAAAGTGTAAAGCCCG |
| 56 | CCAGCAGGCGAAAATCCTGTTTGATGGTGGTCAGAATGCGTAC |
| 57 | CAATCATATCATTATAACCAAAATAATCATGGAGGTTTCTTTGCTCGTCGGA |
| 58 | CTATCAGGTCAACCGTTTTTAGAAATTGCTAGCGGTGC |
| 59 | AGTCTGGAGCAAACAAGAGCGGAGACATGCCTGAG |
| 60 | ATTTTCGCAAGAAAAATAAGGCCCATTAAAAAAAGGGACAT |
| 61 | TGGGGAATGCAGCATTTGGGACAA |
| 62 | GCTGATTGCCGTTCCGCTGCAGCCGAATATAACAATA |
| 63 | ATGGCTTTCCGGCACCTGGTGAAGATATTTAATTTGCGGGTTCAA |
| 64 | GTCCACTAAATCCCTTTAAACCAAGGCGGGCCTGT |
| 65 | CACCAGACGATATCAAAATTATTTGCAATCATTTTCTAAAGCA |
| 66 | TGAAACTCGTTTATAGATTTAGTTAA |
| 67 | ATTCAACTTGCCTAATAGCGGGGT |
| 68 | AGACGATCCAGCGCAGGTTTTCACGGTCATAC |
| 69 | TTAATGAAGATCGTGCCGGAAACCAGGCAAAGCGCCCCC |
| 70 | GGGGTTTCTGCCAGCAGAGTTGCAGAGTAGTA |
| 71 | GTGTATTTTTGAATGGCTAGGAGCACTTATTCATCCTGATTCGCCTCCC |
| 72 | TCGGGAGAAACAATAAAGGAT |
| 73 | AAGGTCGGTCGCTGAGGCTAAGGCACCAACACTATTAATAAATAATGAA |
| 74 | ATATTACCAAATACCTGTAATTCTAAAACTTT |
| 75 | CGCTAAACAACCCTAATTGGAAAAACGCTCATGGGCCAGCCAAGAACAAGGAAAATAG |
| 76 | CTAACTTCTTTGCAGGAGGCCGATTAAAGGGA |
| 77 | TAAACAAAAGAACCTTATGCTTTTCACCAGTGAGACGGGCAACATCACAGTTAGAGG |
| 78 | AGAATTAGATAGAGCCGGCGCTCACAATTCCACA |
| 79 | TTCAAATATCAATATTGAAAAATGCGGAACAAAGAAACCACCAGAAGGTT |

| | |
|---|---|
| 80 | CCGGAACACCGGAACGCTTTGAA |
| 81 | CGGATTCGTTCAATTATTCA |
| 82 | TGTACCAGTAATAATACCGAACGAACCAATCTGG |
| 83 | GTCTCATTTGTTTAATGGAGATAAATTGGCAGATTCACCAGCTTGCCTGTCAGA |
| 84 | CAGTTTCAGCGGAGTGAGATTTTCTGCG |
| 85 | TAGTGAACAAGAAAAATAACCAATCAAACGGG |
| 86 | GTTTCGTCACCAGCGGTTTATAGCATTAATGTTT |
| 87 | AGGTTGAAAATCTCCAAAAACAGCCCTAGGAACGCAGATGGGCCGGTGCG |
| 88 | CAGACCAGGCGCATATTGAGATGAGACTGGTGTTC |
| 89 | AAAGCCCAATAGCCAGTAATTATTTACAACAGAGGTGAGGCGGTGCTG |
| 90 | GAGCACGTATCCAAATTTACCGCGAACTGGCAAAGAAACGCAAAGACATTAC |
| 91 | CCATCGCCGTAAAAAAAGC |
| 92 | AAAACATTTTGTGCAACAGTGCCACGCTGAGAATAACGCGAGGTCCAGAC |
| 93 | CGGATCCGCCGGGCGCGGTATGCCAACAAATCATACGTC |
| 94 | GCGAATCATTTTTAATAAAGCTTTTCACGCCGCTTTT |
| 95 | AGGATTAGAGAAAAATCACTCAAATGTCAGTGAAT |
| 96 | AGCGGGGTTTTGCTCAGTAC |
| 97 | AATCATCTTCTAGCAAGGCAGAATCAAGTTTGCCTCCGCCAGCAAGAAATT |
| 98 | CAAACCATCAATATGATATTC |
| 99 | GCGTTCCAAGATAATCGGCCAGTTTGGAACAAGA |
| 100 | GAAACAAAGTACAGAGATTTA |
| 101 | CCTAGGAATTGAATGATGAACTTTTACA |
| 102 | CATTGACCTAAAAATTACCTTCAAATATAATTTTAAAAGTTTGAGTAACA |
| 103 | GCTCAAAGCGAACCAGACCCAGAAGCCGGAATCGCCAGAACGCAAGCGG |
| 104 | AAATTAGTAATAACATCATCACACGAATTTAGGCACCG |
| 105 | ACCCACCGGAAAAAGAAGGGAAGGTTGAAGTATTAGACTTTACAAACAAAACA |
| 106 | TGATTTCGAGGTGAATTTCCATGTACCTGTGAGCG |
| 107 | GTGTATCTCTGCCAGAATCAGCACGTACAGTGTAGAACGTCAGCGGGCT |
| 108 | GCTGAGACTCCTCAATTAACGGGATCAGAAACAGCGGA |
| 109 | GTAAGGTACTGGTAATAAGTTGAGAAGGATACAAAGG |
| 110 | GCTCATTTGCCGCCAGAGGGTAAAGTTAAACG |
| 111 | CTGGCACTCCAGCCAGCTTTTGATGATAATAAGCAAGGATAGCTTTTTCGTC |
| 112 | CAGCGCCAAGCTCCGTCGGTGGTGCCATGGCGCTTTTATATTTTATACATAATAAA |
| 113 | GGGTGGAATTCGTAGCGAGAGATAACAGTCAGCAGCGTAAAGGAATTGCGAATTTTT |
| 114 | TATAAACAGCCAAAGTCA |
| 115 | GGACGTTGGGAAGAAATAGCCGGAACGTAATG |
| 116 | AGAGTTCGCTATTATTATACTATTAAAGCCAGAATGGAAAGCGCA |
| 117 | TTCGTAAAAGGAAACAGTAGCGACCGGAAACGTCACCAATGGTG |
| 118 | AGCTGTGATAACAAAATTAAGTTGGCACTCGTATT |
| 119 | CCCTGAGAGTATTAAGAG |
| 120 | AAAACGACGGCCAGTTTTCATCAATAGTAGTCAGCTTGCCAACAACCATCGCCCAGA |

| | |
|---|---|
| 121 | GGAAAAAAATTCGTGTACCAGGAACAACAAAGACAGCATCGGAGGAC |
| 122 | TCACCTTCAGTATTATTAATTTATAAAGTACATATAATGATTAAG |
| 123 | AGAAGAATAACCACATAAAAAAATCATAGCAG |
| 124 | ACCGCAGCAAAAGCGCGTTTTCATCGGC |
| 125 | TCGCAATAGTATTATCATCATATTCCTGATTATCTTG |
| 126 | CTGAAAGCGGAGACCTTCATCAAGAGTAATCTTGAGAG |
| 127 | ACGTGGACTCCAACGT |
| 128 | ATACAGTTTCGACAAAATCAACAGTTGAAAAAAC |
| 129 | CACCCAGCAGTGTTTTGAGGCCACCGAGTAAAGCGTAAGAAAGCCAAAAGCCTG |
| 130 | AGAGGTGAATTAGTTTTGCCGAGGATCCACTGGTGTGTTCAGCGGCA |
| 131 | AATCCTGAAGATGATTTAGATTAAGA |
| 132 | TCCTTTACAGAGAGAATAACATAAAAACAGGGTTACGCATTTTATTTAGGT |
| 133 | AGATTTGAAATAGAGGCATTAAGTTTAAATAATAACGGAATACCCTG |
| 134 | TTAATCTAAAATATCTTTATTAGTCTTACTAGAAACGCTCAACGCGA |
| 135 | GACCCCCATTGCGCCGGCGCGAGCGCATCAAT |
| 136 | CGCCTGGCCCTGAGACGCGTGCCATAGCGTCTGCTGTAGTCGAGCTTGATAAATT |
| 137 | GAGCAACCTAAATCCGCGACCTGCTCCATGTTACTAATCTACGTCAT |
| 138 | ATATTCACAAACAAATGTACATCGTTGCTTCTAGACAAAGCATAG |
| 139 | ACGTAAATCGGTCATTAAATTCTTTCCAGTATAGCCCGGAATAG |
| 140 | CAAAGGGCTCAAAAATCAAGCCGTGTATGTTAGCAAAGGTGCTGCGGC |
| 141 | CGCCCCTTCTGACCTGAAAAGAGTCTGTGCGGGAGTATCCTGAAATAATA |
| 142 | AAATATCCCATATGTTCAGGCAAATCCAGTGATGACAGTTGGGCGGTTGTAAATC |
| 143 | ATATCGCGAAGAGGAAAAATGTTTGTTTAATTCCCTTCAC |
| 144 | CTATTATTCTGAAACCCGTATAA |
| 145 | TAAAGTCACCGACCCCTTATTAGCGTTTCAGA |
| 146 | TATATAAATCAAAAGAAT |
| 147 | TTTACGAGCGTGGCGTTTTATAGCCGATTTACCAGCGCCAAAGTATC |
| 148 | TTCCAGATGCCGGGTTACGCAAACGCAACGCAAGAAAACAGGGAGAGATCTTAGGATT |
| 149 | CGCTGTAACCTCCATGCTGATCTAATGCAGAAC |
| 150 | AAAAACGGATAAGCAGCAACCGCAAGATGCGGTATCATTTCGCGTAA |
| 151 | GTAAATGAAATAGAAAAAAACATTGGAAGTTTATCCCTTACCCGGGTA |
| 152 | TCTGTAGCAATAGAACGCGACTTTCCAG |
| 153 | AATTGGGCGGCTGGCTTTGCATCAAAAAGATTTTTTAATCTC |
| 154 | TAATCATTGACAGATGAACGGTGTA |
| 155 | ATAAATATTTTAGACACCGCCACGCTCAATCGTCTGCCTT |
| 156 | GCAACCGACAAATCATCGTAGTAATATCGTCTATCACTTTGAC |
| 157 | TCTACTAACATTAAAGTAACACCACCCTCATTTTCAGGGATAG |
| 158 | ACTCCTTAAAGCGCATATTTTTTGTTTAACGGAAAAACCCAGAACA |
| 159 | CCGAGCTCGTTTTTCGATTTTAAGAACTGGCAGGGAACCCATGAGGA |
| 160 | TTTGGAATTAATTTTCCCTTATATGTAAGGCT |
| 161 | TAATTTCATTAAAGGTGAATACAAAAGGAGTAGGGCTCCCGACTTCCATCAC |

| | |
|---|---|
| 162 | GACGACAAACTCATCGTTGCAACATACGAGCATGTAG |
| 163 | GAAAGGCAATCGATGGTAAAACACAGTTAA |
| 164 | TATCACCAGTAGCACCACCACG |
| 165 | CGTCGGATGTTGGGTAACGCCA |
| 166 | AAATCCTTTGCCCGAATTTCAGGTTAACAATTAGACTG |
| 167 | GTCACCTCACCGGAAACAATCG |

**Table S3.** Closing staples and fluorescently labelled staples used in Version 1 of the nanosensor. For the closing interactions, the number of the caDNAno helix is indicated (see Fig. S17). Yellow and Red indicate complementary sequences. Green indicates the toehold used for opening. Blue indicates additional spacer elements.

| **Name-Helix ID** | **Sequence (5'->3')** |
|---|---|
| Row1-top16 | ATACATCTATTATATATTGAGGGTAATTGAGCGCTGAAACCGATATC CGGTAACA |
| Row1-top13 | ATACATCTATTATATATTCCACCACCAGAGCCGTTAGC |
| Row1-bot24 | AATAGATGTATTTATATATTGCGGGACGTTGGGAAGAAATAGCCGG AACGTAATG |
| Row1-bot29 | AATAGATGTATTTATATATTTTGGGAAGGGCGATGCATCGTAA |
| Row2-top13 | ATACATCTATTATATATTTTCAAGCCTAATTTGCCAGTGAGCTA |
| Row2-top16 | ATACATCTATTATATATTTACCCTCATTTTCGCAAGAAAAATAAGGC CCATTAAAAAAAGGGACAT |
| Row2-bot24 | AATAGATGTATTTATATATTTTCGGCCAACGCGCGGGGAGAG |
| Row2-bot29 | AATAGATGTATTTATATATTTGGCCTCTTCGCTATTACCCACGGGAAT AATTCGCAGGCAAGCCAA |
| Row3-top13 | ATACATCTATTATATATTTTATTGAGTAAGCAGAGCGAACCTTAATT GAGATAGAAGAACTGATAGC |
| Row3-top16 | ATACATCTATTATATATTTTACCTCGGGAGAAACAATAAAGGAT |
| Row3-bot24 | AATAGATGTATTTATATATTTTCGTGCCCAACATACGAGGCATAAAA TGGTCCGATATATAGCCTTTAATTGTATTA |
| Row3-bot29 | AATAGATGTATTTATATATTTTCTGAATGGGATAGGTCACGCAGAAC CGTGTA |
| Row4-top13 | ATACATCTATTATATATTTTTAGAGCAAGAAACAATGAGCTATCTAG CCTTAAACCAGTATAATAC |
| Row4-top16 | ATACATCTATTATATATTTTTTCAGAGCCGCCACCCTGCCATCTTCCT GAGCATCATAATTTAATG |
| Row4-bot24 | AATAGATGTATTTATATATTTTTCTTGAAAGATTCATCAGTTACGGAG ATCATCTTT |
| Row4-bot29 | AATAGATGTATTTATATATTTTTTGCAAGCGGCCAGTTTTCATCAATA GTAGTCAGCTTGCCAACAACCATCGCCCAGA |
| Green_dye_non_hybridizing | AATTCTCAGAGCAATTCTGATGGGTAATCATAGCTGTTTCCTGTG- [ATTO542] |
| Red_dye_non_hybridizing | GTTAGAATCAGAGCGGTACAAAATAGAAGGCTGGAAAC- [ATTO647N] |
| Red_dye_1nt_spacer | GTTAGAATCAGAGCGGTACAAAATAGAAGGCTGGAAACTCTGACGG C-[ATTO647N] |
| Green_dye_1nt_spacer | AATTCTCAGAGCAATTCTGATGGGTAATCATAGCTGTTTCCTGTGTG CCGTCAG-[ATTO542] |

**Table S4.** Modified (closing, stabilization and fluorescently labelled) staples used in Version 2 of the sensor. For the closing interactions, the number of the caDNAno helix is indicated (see Fig. S17). Yellow and Red indicate complementary sequences. Green indicates the toehold used for opening. Blue indicates additional spacer elements.

| **Name-Helix ID** | **Sequence (5'->3')** |
|---|---|
| Row1-top13 | ATATCACGAGCACATCAGTACGAGGGTAATTGAGCGCTGAAACCGATATCC GGTAACA |
| Row1-top16 | GTGCTCGTGATATTTTTCCACCACCAGAGCCGTTAGC |
| Row1-bot24 | GTGCTCGTGATATTTTTGCGGTTTGCGTATTGGGCGCCA |
| Row1-bot29 | ATATCACGAGCACATCAGTACTTGGGAAGGGCGATGCATCGTAA |
| Row2-top13 | ATATCACGAGCACATCAGTACTCAGAGAGATAACCCACAAGA |
| Row2-top16 | GTGCTCGTGATATTTTTTTTTACCCTCAGAGCCGCAGATGAAT |
| Row2-bot24 | GTGCTCGTGATATTTTTTTTTTCGGCCAACGCGCGGGGAGAG |
| Row2-bot29 | ATATCACGAGCACATCAGTACGGCCTCTTCGCTATTACCCACGGGAATAATT CGCAGGCAAGCCAA |
| Row3-top13 | ATATCACGAGCACATCAGTACTTATTGAGTTAAGCCCATCTTACCAACGCTA TAGAC |
| Row3-top16 | GTGCTCGTGATATTTTTTTTTTTACCGCCACCCTCAGAGCCACC |
| Row3-bot24 | GTGCTCGTGATATTTTTTTTTTTCGTGCCAGCTGCATACGAACTA |
| Row3-bot29 | ATATCACGAGCACATCAGTACTTCTGGCGAAAGGGGGATGTGC |
| Row4-top13 | ATATCACGAGCACATCAGTACTTTAGAGCAAGAAACAATGAGCTATCTAGC CTTAAACCAGTATAATAC |
| Row4-top16 | GTGCTCGTGATATTTTTTTTTTTTTTCAGAGCCGCCACCCTGCCATCTTCCTG AGCATCATAATTTAATG |
| Row4-bot24 | GTGCTCGTGATATTTTTTTTTTTTTTCTTTCCAGTCGGGAAACCTGT |
| Row4-bot29 | ATATCACGAGCACATCAGTACTTTTGCAAGGCGATTAATCTCCGTGGGAAC AAAGAGCCACTGA |
| 5bp_stabilisation_1 | CATTGCACCCACGCAACCAGCTTGACGTTGTTTTTTAGACC |
| 5bp_stabilisation_2 | TTAATTGCGTTGCGCTCACTGCCTTTTTCACAC |
| 5bp_stabilisation_3 | AACGGTACGCCAGAATCCTGAGATACAATTTGTTTTGATACCGATTTTTGTG TG |
| 5bp_stabilisation_4 | TTGAGGGAATATTGACGGAAATTACAAAATCATTTTTGGTCT |
| Red_dye_3nt_spacer | GTTAGAATCAGAGCGGTACAAAATAGAAGGCTGGAAACTTTCTGACGGC- [ATTO647N] |
| Green_dye_3nt_spacer | AATTCTCAGAGCAATTCTGATGGGTAATCATAGCTGTTTCCTGTGTTTGCCG TCAG-[ATTO542] |

**Table S5.** Biotinylated staples for immobilization of DNA origami nanostructures used in both versions of the sensor. Blue indicates additional spacer elements.

| Name | Sequence (5'->3') |
|---|---|
| Biotin1 | 5'Biotin-TTTATTGCCGGTTGATAGTCAGTGCCTTGAGTAACAGTGCATGAA |
| Biotin2 | 5'Biotin-TTTATTATAGTGGAAGCAATAGAGCTTACCCTCATATATTTTTGTAC |
| Biotin3 | 5'Biotin-TTTCAAACTCTTCCTGTTGACCGT |
| Biotin4 | 5'Biotin-TTTATACACCGCCAAAATACAGGTA |

**Table S6.** DNA-DNA closing interactions used for XhoI nanosensors. Shown in purple is restriction site of XhoI. The number of the caDNAno helix is indicated (see Fig. S17).

| Name-Helix ID | Sequence (5'->3') |
|---|---|
| Row1-top 13 | ATACTCGAGTTATATATTGAGGGTAATTGAGCGCTGAAACCGATATCCGGTAACA |
| Row1-top16 | ATACTCGAGTTATATATT CCACCACCAGAGCCGTTAGC |
| Row1-bot24 | AACTCGAGTATTTATATATTGCGGGACGTTGGGAAGAAATAGCCGGAACGTAATG |
| Row1-bot29 | AACTCGAGTATTTATATATT TTGGGAAGGGCGATGCATCGTAA |
| Row2-top 13 | ATACTCGAGTTATATATTTTCAAGCCTAATTTGCCAGTGAGCTA |
| Row2-top16 | ATACTCGAGTTATATATTTACCCTCATTTTCGCAAGAAAAATAAGGCCCATTAAAAAAAGGGACAT |
| Row2-bot24 | AACTCGAGTATTTATATATTTTCGGCCAACGCGCGGGGAGAG |
| Row2-bot29 | AACTCGAGTATTTATATATTTGGCCTCTTCGCTATTACCCACGGGAATAATTCGCAGGCAAGCCAA |
| Row3-top 13 | ATACTCGAGTTATATATTTTATTGAGTAAGCAGAGCGAACCTTAATTGAGATAGAAGAACTGATAGC |
| Row3-top16 | ATACTCGAGTTATATATTTTACCTCGGGAGAAACAATAAAGGAT |
| Row3-bot24 | AACTCGAGTATTTATATATTTTCGTGCCCAACATACGAGGCATAAAATGGTCCGATATATAGCCTTTAATTGTATTA |
| Row3-bot29 | AACTCGAGTATTTATATATTTTCTGAATGGGATAGGTCACGCAGAACCGTGTA |
| Row4-top 13 | ATACTCGAGTTATATATTTTTCAGAGCCGCCACCCTGCCATCTTCCTGAGCATCATAATTTAATG |
| Row4-top16 | ATACTCGAGTTATATATTTTTTCAGAGCCGCCACCCTGCCATCTTCCTGAGCATCATAATTTAATG |
| Row4-bot24 | AACTCGAGTATTTATATATTTTTCTTGAAAGATTCATCAGTTACGGAGATCATCTTT |
| Row4-bot29 | AACTCGAGTATTTATATATTTTTTGCAAGCGGCCAGTTTTCATCAATAGTAGTCAGCTTGCCAACAACCATCGCCCAGA |

**Table S7.** Antigen modified staples that were used in a sensor for anti-Dig and anti-DNP antibodies. The number of the caDNAno helix is indicated (see Fig. S17). Blue indicates additional spacer element.

| Name-Helix ID | Sequence (5'->3') |
|---|---|
| Row1-top16 | [DNP]TGAGGGTAATTGAGCGCTGAAACCGATATCCGGTAACA |
| Row1-top13 | [DNP]TCCACCACCAGAGCCGTTAGC |
| Row1-bot24 | [DNP]TGCGGGACGTTGGGAAGAAATAGCCGGAACGTAATG |
| Row1-bot29 | [DNP]TTTGGGAAGGGCGATGCATCGTAA |
| Row4-top13 | [DIG]TTTAGAGCAAGAAACAATGAGCTATCTAGCCTTAAACCAGTATAATAC |
| Row4-top16 | [DIG]TTTTCAGAGCCGCCACCCTGCCATCTTCCTGAGCATCATAATTTAATG |
| Row4-bot24 | [DIG]TTTCTTGAAAGATTCATCAGTTACGGAGATCATCTTT |
| Row4-bot29 | [DIG]TTTTGCAAGCGGCCAGTTTTCATCAATAGTAGTCAGCTTGCCAACAACCA TCGCCCAGA |

**Table S8.** Sequences of opening strands used in different sensor constructs. Red indicates the mismatch.

| Name | Sequence (5'->3') |
|---|---|
| 15 bp interaction | ACTGATGTGCTCGTG |
| 17 bp interaction | ATATATAATAGATGTAT |
| 17 bp mismatch GC | ATATATAATA**C**ATGTAT |
| 17 bp mismatch AT | ATATATAATAG**T**TGTAT |
| 17 bp mismatch GC+AT | ATATATAATA**CT**TGTAT |
| 17 bp mismatch Toehold AT | AT**T**TATAATAGATGTAT |
| 17 bp mismatch Toehold TA | ATA**A**ATAATAGATGTAT |

**Table S9.** Fit results as well as buffer compositions for the titration data shown in Fig. 2.

| **Sample** | **Buffer** | $\mathbf{K_{1/2}}$ | $\mathbf{n_H}$ |
|---|---|---|---|
| 2x13 | 10 mM Tris, 10 mM $MgCl_2$, 50 mM NaCl | 100.08 ± 10.43 nM | 0.9782 ± 0.084 |
| 2x13 | 10 mM Tris, 10 mM $MgCl_2$, 200 mM NaCl | 195.05 ± 12.36 nM | 0.8709 ± 0.041 |
| 2x13 | 10 mM Tris, 10 mM $MgCl_2$, 400 mM NaCl | 601.19 ± 19.57 nM | 0.8075 ± 0.019 |
| 2x13 stabilized | 10 mM Tris, 10 mM $MgCl_2$, 50 mM NaCl | 704.97 ± 64.84 nM | 0.7997 ± 0.049 |
| 4x13 | 10 mM Tris, 10 mM $MgCl_2$, 50 mM NaCl | 1087.31 ± 85.60 nM | 1.5459 ± 0.1524 |
| 6x13 | 10 mM Tris, 10 mM $MgCl_2$, 50 mM NaCl | 2025.14 ± 102.35 nM | 1.7335 ± 0.1255 |
